# Flexibility of co-evolutionary patterns in ectoparasite populations: genetic structure and diversity in *Apodemus* mice and their lice

**DOI:** 10.1101/065060

**Authors:** J. Martinů, V. Hypša, J. Štefka

**Affiliations:** Biology Centre CAS, České Budějovice, Czech Republic; Faculty of Science, University of South Bohemia, České Budějovice, Czech Republic

**Keywords:** co-evolution, dispersal, genetic diversity, isolation by distance, Apodemus, Polyplax

## Abstract

Host-parasite co-evolution belongs among the major processes governing evolution of biodiversity on the global scale. Numerous studies performed at inter-specific level revealed variety of patterns from strict co-speciation to lack of co-divergence and frequent host-switching, even in species tightly linked to their hosts. To explain these observations and formulate ecological hypotheses, we need to acquire better understanding to parasites’ population genetics and dynamics, and their main determinants. Here, we analyse the impact of co-evolutionary processes on genetic diversity and structure of parasite populations, using a model composed of the louse *Polyplax serrata* and its hosts, mice of the genus *Apodemus*, collected from several dozens of localities across Europe. We use mitochondrial DNA sequences and microsatellite data to describe the level of genealogical congruence between hosts and parasites and to assess genetic diversity of the populations. We also explore links between the genetic assignment of the parasite and its host affiliation, and test the prediction that populations of the parasite possessing narrower host specificity show deeper pattern of population structure and lower level of genetic diversity as a result of limited dispersal and smaller effective population size. We demonstrate an overall complexity of the co-evolutionary processes and their variability even among closely related lineages of the parasites. In the analysis of several sympatric parasite populations, we find strong evidence for the link between the width of host specificity and genetic diversity of parasites.

## Introduction

Genetic structure and diversity of populations, and their genetic connectivity are the key elements in long-term population survival and evolution, and in the origin of new species. Formation of the genetic structure is contingent on an interplay of various factors, such as the environment, life strategy, population history, etc. Despite the fact that parasites represent one of the most common ecological strategies (Price 1980), majority of our knowledge on the population processes generating genetic diversity is derived from the studies on free-living organisms. The generally accepted view holds that species occupying large interconnected habitats tend to possess larger, more diverse populations, whereas species with isolated populations and/or recently bottlenecked species show reduced diversity (Allendorf *et al.* 2013). However, even in continental, highly mobile species, the level of local genetic diversity may differ and gene flow between the populations may be affected by moderate environmental differences (Lemoine *et al.* 2016).

In parasites, particularly in those with life-cycles closely bound to their hosts, as for example parasitic lice, the host represents the sole parasite’s environment. In such cases, the parasites typically develop a strong narrow host specificity, and their population structure, diversity, and speciation processes are assumed to be strongly, or even entirely, determined by their host. At an inter-specific, phylogenetic level, this results in a parallel evolution, which may lead to an almost perfect fit between the host’s and parasite’s phylogenies (Hughes *et al.* 2007; Light & Hafner, 2008). In most cases, however, host switches blur the co-evolutionary signal, even in highly host-specific parasites (Ricklefs *et al.* 2004; Banks *et al.* 2006). For example, in the sucking lice *Polyplax arvicanthis* and mice of the genus *Rhabdomys* in South Africa, du Toit *et al.* (2013) found that two sympatric lineages of *Polyplax arvicanthis* showed only limited congruencies with their hosts.

Possible processes causing phylogenetic inconguencies between the host and parasite have often been discussed in parasitological literature, and a complex conceptual background has been developed (Page 2003; Clayton *et al.* 2004; Toon & Hughes 2008; Lion & Gandon 2015). For example, biogeography, social behaviour and vagility of the hosts were suggested to affect the level of congruence in host-parasite equally or even in greater extent than the bionomy and life history traits of the parasite. However, estimating the degree of intimacy for a particular host-parasite association is not a simple task. It may be even counter-intuitive, if previously unforeseen factors are involved in the interaction. This was for instance illustrated by Engelbrecht *et al.* (2016) in their study on a temporary parasite *Laelaps giganteus*, where authors found significant co-diversification pattern between mites and *Rhabdomys* mice, even though *Laelaps* mites spend most of their life off the host in their nests (Mullen & O’Connor 2002). Among the reasons, why the *Laelaps* mites show seemingly higher level of intimacy than the permanent *Polyplax* lice, could be the limited dispersal abilities due to the low abundance and prevalence on the hosts. From similar studies, it becomes obvious, that the key to understanding the co-evolutionary pattern is the investigation of the parasites’ population genetics and dynamics, and their main determinants. At this intra-specific level, the current research showed that parasite diversity and population structure is affected by several factors, mainly shared demographic history (e.g. Nieberging *et al.* 2004; Štefka *et al.* 2011), host dispersal capabilities affecting parasite’s gene flow (e.g. McCoy *et al.* 2003; Štefka *et al.* 2009; van Schaik *et al.* 2014), and the spectrum of parasitized hosts (e.g. Barrett *et al.* 2008; Archie &Ezenwa 2011). Nadler (1995) stressed the role of host specificity, predicting that multihost parasites display shallower population structure due to better chance to disperse.

To our knowledge, only few studies on natural populations of parasites have been designed to allow for addressing these issues, for example the co-evolutionary reconstruction of feather lice species with extremely different host specificities (Johnson *et al.* 2002) or the investigation of two generalist pinworms from reptiles from the Caribbean area (Falk & Perkins 2013). They support the Nadler’s prediction in a general sense, by showing that the parasites with stronger host specificity possessed more pronounced genetic structure. However, while in free-living organisms the effect of the ecological parameters and their shifts on population genetics is well explored (e.g. Lemoine 2016), the extent to which even moderate changes of host specificity shape the structure and genetic diversity of parasites remains largely unknown.

In this study, we address the impact of host specificity on the genetics of parasite populations using the system of a sucking louse *Polyplax serrata* and its hosts, mice of the genus *Apodemus.* Compared to the *Rhabdomys* mice, with parapatric species inhabitting differentiated bioms, (du Toit *et al.* 2013), the *Apodemus* model possesses different geographic and ecological structure. Two most widespread species, *Apodemus flavicollis* and *A. sylvaticus*, co-occur throughout majority of their European distribution in sympatry or even syntopy (Michaux *et al.* 2005). In the eastern part of their geographical distribution they overlap with two other species, *A. uralensis* and *agrarius*, which expanded to central Palearctic from the east quite recently (Suzuki *et al.* 2008). The sister species *A. sylvaticus* and *flavicollis* separated more than 4 million years ago (mya) (Michaux & Pasquier 1974) and responded differently to the Quaternary climatic oscillations. The more adaptable *A. sylvaticus* persisted glaciation in the Iberian peninsula and colonized Europe mainly from there. Greater forest dweller, *A. flavicollis*, did not survive in Iberia and its refugium was connected with the area of Italy and the Balkans (Michaux *et al.* 2005). *A. sylvaticus* occured in the Balkan region during glaciation too, but suffered a genetic bottleneck there (Michaux *et al.* 2003), potentially as a consequence of competition with *A. flavicollis*, which reaches higher abundance when the two species are in sympatry (Michaux *et al.* 2005). The nonuniform evolutionary history of the two species had also impact on the genealogies of their parasites, e.g endoparasitic helminths (Nieberding et al 2004; 2005) and the permanent host-specific parasites, like the sucking lice of the genus *Polyplax*.

The basic genetic structure of *Polyplax*/*Apodemus* system, as revealed by Štefka & Hypša (2008), shows this system as a useful model for co-evolutionary studies at population level, and rises several interesting questions/hypothesis to be addressed in the present study. At the general level, genealogy and current distribution of the lice was clearly coupled with the evolutionary history and distribution of *Apodemus*. However, the specificity and phylogeographical patterns varied across three main mtDNA-based lineages of the parasite (designated as A, B and C). Two lineages, A and B, were more ubiquitous in their distribution and occurred in sympatry, but differed in the degree of their host specificities. Both clades shared *A. flavicollis* as a common host, and mostly occupied sympatric localities in central Europe, but the lineage A also parasitized another species, *A. sylvaticus*, and was found also in western Europe (France and United Kingdom). The lice of the lineage C inhabited mainly *A. agrarius* and *A. uralensis* occurring in the central and eastern areas of Europe. In the present study, we analyze an extensive sample across multiple European countries to answer the following questions: 1. Do the mtDNA *Polyplax* lineages revealed by Štefka & Hypša (2008) retain their integrity and host specificity if analysed with multi-locus data on considerably extended geographical sampling? 2. Do lineages of the parasite show a strict pattern of co-divergence with their hosts, i.e. do genealogies of *Polyplax* lineages correspond with those of their principal hosts, *A. sylvaticus* and *flavicollis*? 3. Is host dispersal the sole determining factor of the parasite gene flow, i.e. do parasites possess similar or even stronger pattern of population structure compared to their hosts? 4. Do the parasitic lineages A and B with different width of host-specificity, follow the Nadler’s rule (Nadler 1995) in the sense of i) deeper population structure in the more host specific lineage, caused by lower dispersal possibilities and ii) significant differences in genetic diversity between sympatric populations correspondingly to the width of their host spectrum?

## Materials and methods

### Host sampling and DNA isolation

A total of 2352 specimens of *Apodemus* hosts were collected across 14 European countries during years 2005-2015. Mice were captured in wooden snap traps (except for few Slovakian samples caught in live traps of Sherman type) and euthanized with ether. *Apodemus* tissue samples (ear or finger tips) were preserved in ethanol and mice were examined for lice by combing the fur with brush. Collected lice were stored in 100% ethanol in the freezer. Field studies were carried out with permits provided by the Czech Republic/European Union or collaborating institutions (Permit Numbers KUJCK 11134/2010 OZZL/2/Ou and 27873/ENV/11); the protocol was approved by the Committee on the Ethics of Animal Experiments of the University of South Bohemia and by the Ministry of the Environment of the Czech Republic (Permit Numbers 13841-11 and 22395/2014-MZE-17214). DNA extractions of individual specimens of lice were performed with QIAamp DNA Micro Kit (Qiagen) into 30µl of AE buffer. Louse skeletons were preserved in 70% ethanol as vouchers. Host DNA was isolated from the host tissue with DNeasy Blood & Tissue Kit (Qiagen) according to manufacturer’s instructions.

### DNA sequencing and population analysis

Fragment of the mitochondrial cytochrome oxidase subunit I gene (COI, 381bp) was amplified for 430 specimen of *Polyplax serrata* lice from 348 *Apodemus* hosts using primers L6625 and H7005 (Hafner *et al.* 1994). These primers, reliably amplifying louse DNA samples of varying quantity and quality, were selected to provide a gross picture of population structure across the whole sample set. For better understanding of the relationships between main mtDNA lineages of lice, a longer fragment of COI (1027bp), together with three nuclear genes VATP21 (304bp), hyp (380bp) and TMEDE6 (215bp), were obtained for selected specimens of *P. serrata* (n=25), using COI primers LCO1490 and H7005 (Folmer *et al.* 1994) and nuclear primers published by Sweet *et al.* (2014). Description of the PCR reactions, thermal cycling conditions and sequencing are provided in Document S1 (Supporting information). Mitochondrial D-loop region with the entire tRNA^Thr^, tRNA^Pro^ and the beginning of the 12S tRNA (1002bp) was gained for 230 individuals of *A. flavicollis* and 93 specimen *A. sylvaticus* with primers 1, 2bis, 3 and 4 (Bellinvia 2004) using PCR conditions described in Document S1 (Supporting information).

Sequences of *Apodemus* and *Polyplax* were assembled in GENEIOUS8.0.2 (Biomatters, Ltd), collapsed into haplotypes using ALTER (Glez-Peña *et al.* 2010), and phylogenies were reconstructed with methods of maximum likelihood (ML) and Bayesian inference (BI). For all analyses the best-fit models (listed in Document S1, Supporting information) were selected according to a corrected Akaike information criterion (AIC) using jModelTest2 (Darriba *et al.* 2012). *P. spinulosa* was used as outgroup in the phylogeny of the parasite. *A. sylvaticus* and *flavicollis* phylogenies were rooted with 3 individuals of the other species (tree of *A. sylvaticus* with *A. flavicollis* and vice versa). Bayesian topologies were conducted in MrBayes 3.2.4 (Ronquist *et al.* 2012) and all analyses consisted of two parallel Markov Chain Monte Carlo simulations with 4 chains 10 million generations long and sampling frequency of 1000. Convergence of parameter estimates and their ESS values were checked in software TRACER 1.6 (Rambaut *et al.* 2014). 2.5 million generations (25%) were discarded as burn-in. Maximum likelihood analyses were computed using PhyML 3.0 (Guidon *et al.* 2010) with 1000 bootstrap replicates to obtain nodal support.

Haplotype networks for the parasite and host datasets, and a 95% parsimonious connection limit, were reconstructed with statistical parsimony software TCS Networks (Clement *et al.* 2002) implemented in the package PopART (http://popart.otago.ac.nz). Standard diversity indices (nucleotide diversity, haplotype diversity, Theta estimates of mitochondrial effective population size) were computed in DNASP 5.10.1 (Librado & Rozas 2009) for each mitochondrial DNA (mtDNA) clade of the parasites and hosts, and for geographical regions that served as potential refugia for one of the host species compared with the rest of the Europe (Fig. S1, Supporting information). To reveal recent demographic changes within geographic regions or clades, Tajima´s D, Fu and Li´s D*, Fu and Li´s F*, Fu´s Fs and R_2_ statistic tests were calculated. Hierarchical analyses of molecular variance (AMOVA; Excoffier *et al.* 1992) using ϕ indexes were calculated for both parasitic and host mtDNA datasets divided with respect to mtDNA clades or geographic regions as described above (Fig. S1, Supporting information). AMOVA analyses were calculated in ARLEQUIN 3.5.2.2 (Excofier & Lischer 2010), significance was tested with 10 000 permutations.

### Microsatellite genotyping and population structure

To explore recent genealogical processes within species and between populations of both parasites and their hosts, microsatellite loci were incorporated into the study. For 458 individuals of *Polyplax serrata* contained in the mtDNA analysis, 16 microsatellite loci were amplified in 4 multiplex PCR assays using primers and PCR multiplexes developed by Martinů *et al.* (2015). All microsatellite loci were tested for departure from Hardy–Weinberg equilibrium (HWE) and linkage disequilibrium (LD) between loci pairs for all populations in GenAlEx 6.5 (Peakall & Smouse 2012). To determine whether populations of lineages A, B, C and from Baikal (referred to hereafter as S, N, Aa and Ape, respectively) form distinct clusters, as in mtDNA phylogenies, or whether they admixed, several approaches of population clustering were used. First, multivariate technique of Principal Coordinate Analysis (PCoA) was computed from genetic distance matrix calculated across multiple loci for each pair of individuals. The same analysis was performed also on the level of populations. PCoA analyses together with an assignment test of S and N lineages were performed in GenAlEx 6.5 (Peakall & Smouse 2012). Then, in POPTREEW (Takezaki *et al.* 2014) pairwise DA values (Nei *et al.* 1983) were calculated and neighbour-joining (NJ; Saitou & Nei 1987) tree was built with 1000 bootstrap replications. Finally, two Bayesian methods based on distinct computational algorithms, Structure 2.3.4 (Pritchard *et al.* 2000; Falush *et al.* 2007; Hubisz *et al.* 2009) and BAPS 6.0 (Corander *et al.* 2008), were used to explore population structure in the whole *Polyplax* dataset and then separately in the S and N lineages (settings provided in Document S1, Supporting information).

For *A. flavicollis* and *sylvaticus* 7 microsatellite loci were amplified in two multiplex assays, following Harr *et al.* (2000) and Aurelle *et al.* (2010). Additional 5 loci specific only for *A. flavicollis*, using multiplexes according to Aurelle *et al.* (2010), and 10 loci specific for *A. sylvaticus* (Makova *et al.* 1998; Harr *et al.* 2000) were amplified to complement datasets of each species. Altogether 230 individuals of *A. flavicollis* and 93 individuals of *A. sylvaticus* were genotyped and all sampled specimens were included in mtDNA phylogenies as well. All loci were tested for departure from HWE and for LD between pairs of loci in GenAlEx 6.5 (Peakall & Smouse 2012).

Same analyses as described above for *Polyplax* lice were done for both *Apodemus* species to reconstruct their population structure and to reveal the level of integrity or mixing of individual mtDNA lineages within and between populations. Shared loci for *A. sylvaticus* and *flavicollis* (7) were analysed with Bayesian approaches (Structure, BAPS), then distance- based methods (PCoA on population and individual level, POPTREEW with 1000 bootstraps) and assignment test were calculated to confirm integrity of the two species. On the intra-specific level Bayesian methods with K=1-20 and the same settings as for *Polyplax* were used (Document S1, Supporting information). PCoA of individuals and populations and NJ phylogeny using DA distances (in POPTREEW) were done for completeness of the picture of genetic differentiation between populations.

### Distribution of genetic diversity in *Polyplax* and *Apodemus*

For *Polyplax* populations from S and N lineages, and for both *Apodemus* species, whose species identity was confirmed by population clustering methods (BAPS, Structure and PCoA analyses), pairwise *F*_*ST*_ were calculated, with 9999 permutations to test significance of the results. In cases of high mutation rates of microsatellite loci, *F*_*ST*_ type estimates are known to be biased towards low values instead of reflecting the true level of genetic differentiation (see Jost 2008). We used R package diveRsity (Keenan 2013) to visualize the effect of possible bias in our *F*_*ST*_ results by comparing the relationship between polymorphism (mean number of alleles per locus) and differentiation (calculated for *F*_*ST*_, *G*_*ST*_, *G*_′*ST*_ and *D*_*JOST*_).

To assess the influence of geographic distance on genetic relatedness, Mantel tests (Mantel 1967) were used to test for isolation by distance (IBD) using microsatellite estimates of genetic differentiation (either *F*_*ST*_, D_JOST_ or G_ST_) and geographic distances separately for both *Polyplax* lineages and both *Apodemus* species in the R package adegenet (Jombart 2008). Statistical significance was computed by 10 000 random permutations. Because the effect of IBD could play different role at different geographic scales, we analysed the spatial autocorrelation coefficient (*r*) for *Polyplax* S and N lineages and both *Apodemus* hosts. The analyses were performed in GenAlEx 6.5 (Peakall & Smouse 2012), where *r* was calculated for increasing distance classes with 95% confidence interval obtained by 1000 bootstrap replicates and 10 000 permuted *r* values (Smouse & Peakall 1999; Peakall *et al.* 2003).

The impact of host genealogy on genetic structure of the parasite was evaluated by correlating F_ST_ (and G_ST_) matrixes of each of the *Polyplax* lineages and its host species using Mantel tests in R package adegenet and GenAlEx6.5.

To determine the plausible difference in the depth of genetic structure in microsatellites among S and N mtDNA clades of *Polyplax*, *F*_*ST*_ and Gene diversity (*H*) indices were calculated in FSTAT 2.9.3.2 (Goudet 1995; 2001) with P-values determined by 10 000 permutations. Genetic indices were calculated for seven localities scattered in five European countries, from which sufficient sample for both sympatric lineages was obtained. Analysis was performed in GenAlEx 6.5 (Peakall & Smouse 2012).

## Results

### Phylogeny and genealogy of *Polyplax serrata* lice

Partial COI genes were sequenced for 430 louse specimens and aligned with 126 sequences obtained by Hypša and Štefka (2008). Final mitochondrial dataset contained sequences of 556 *Polyplax* specimens (Fig. 1, Table 1).

**Fig. 1:**
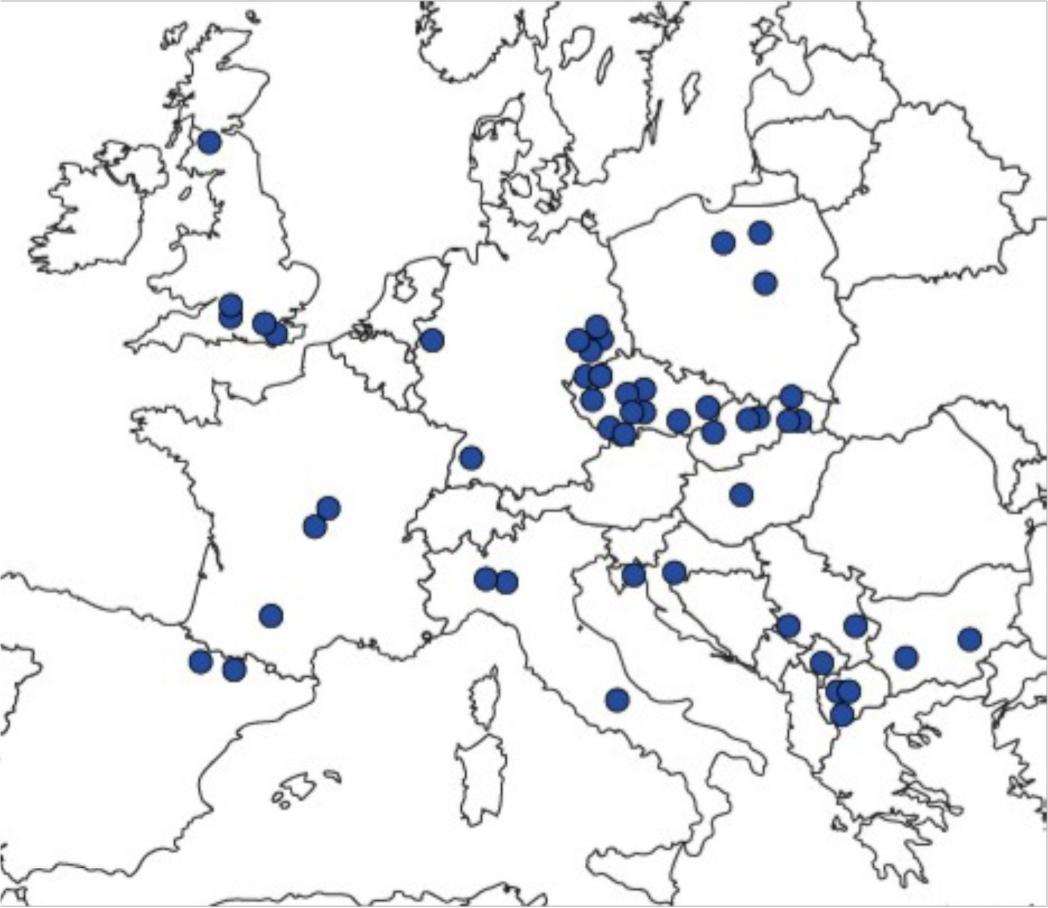
Map of European sampling localities. Collection site of the single Asian locality (Baikal Lake) not shown.

**Table 1:**
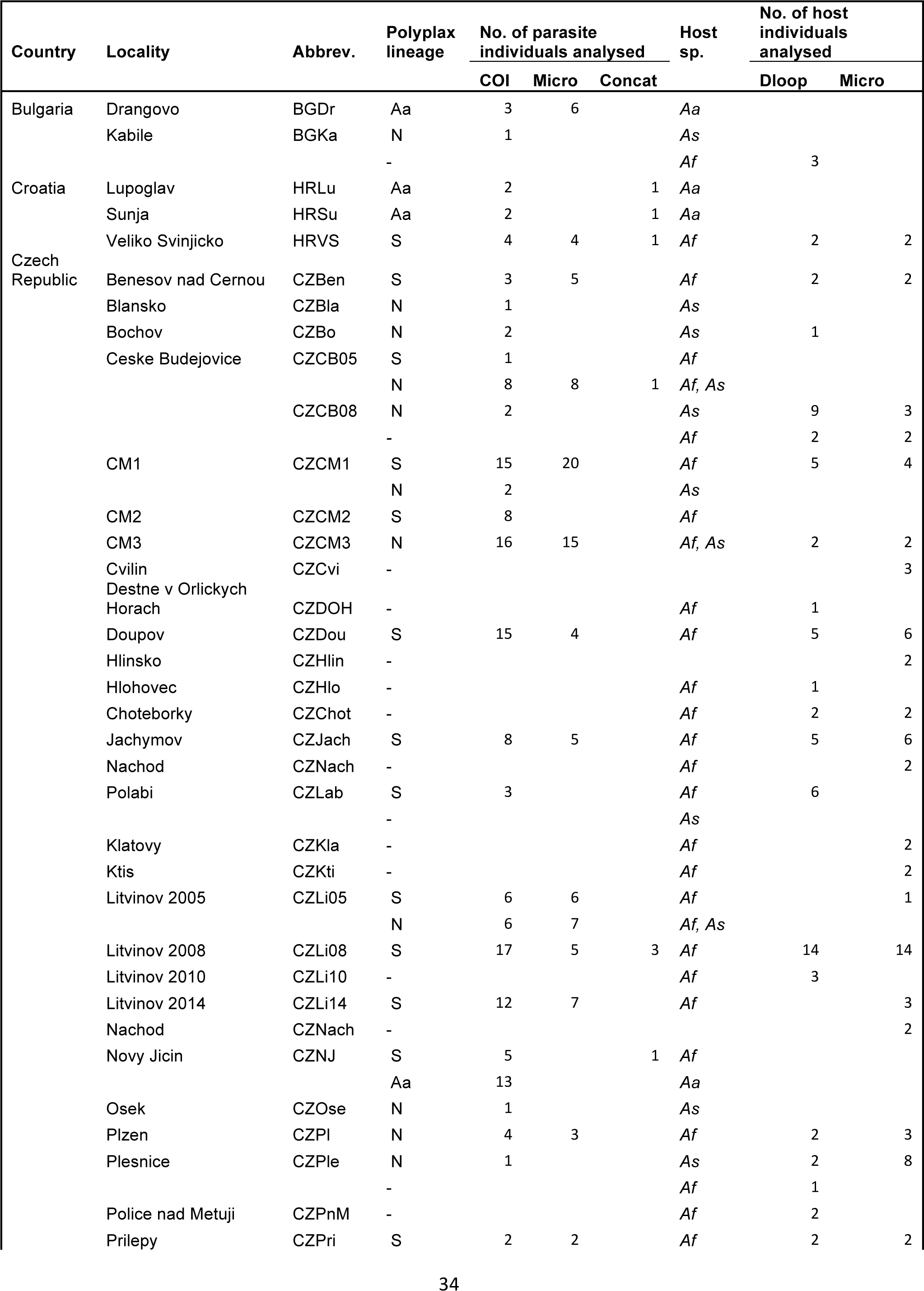
List of sampling localities providing numbers of samples analysed for each organism and marker.

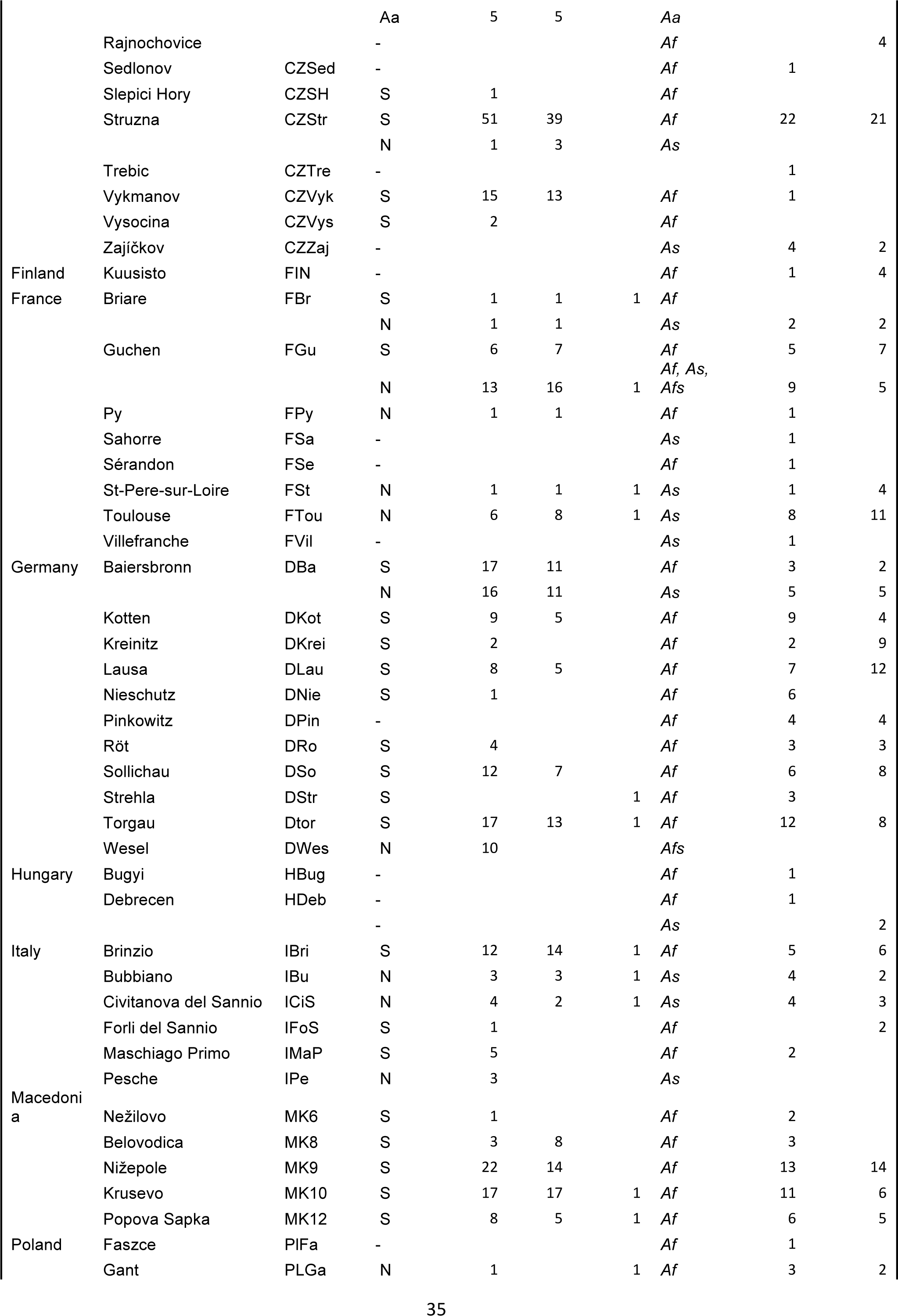

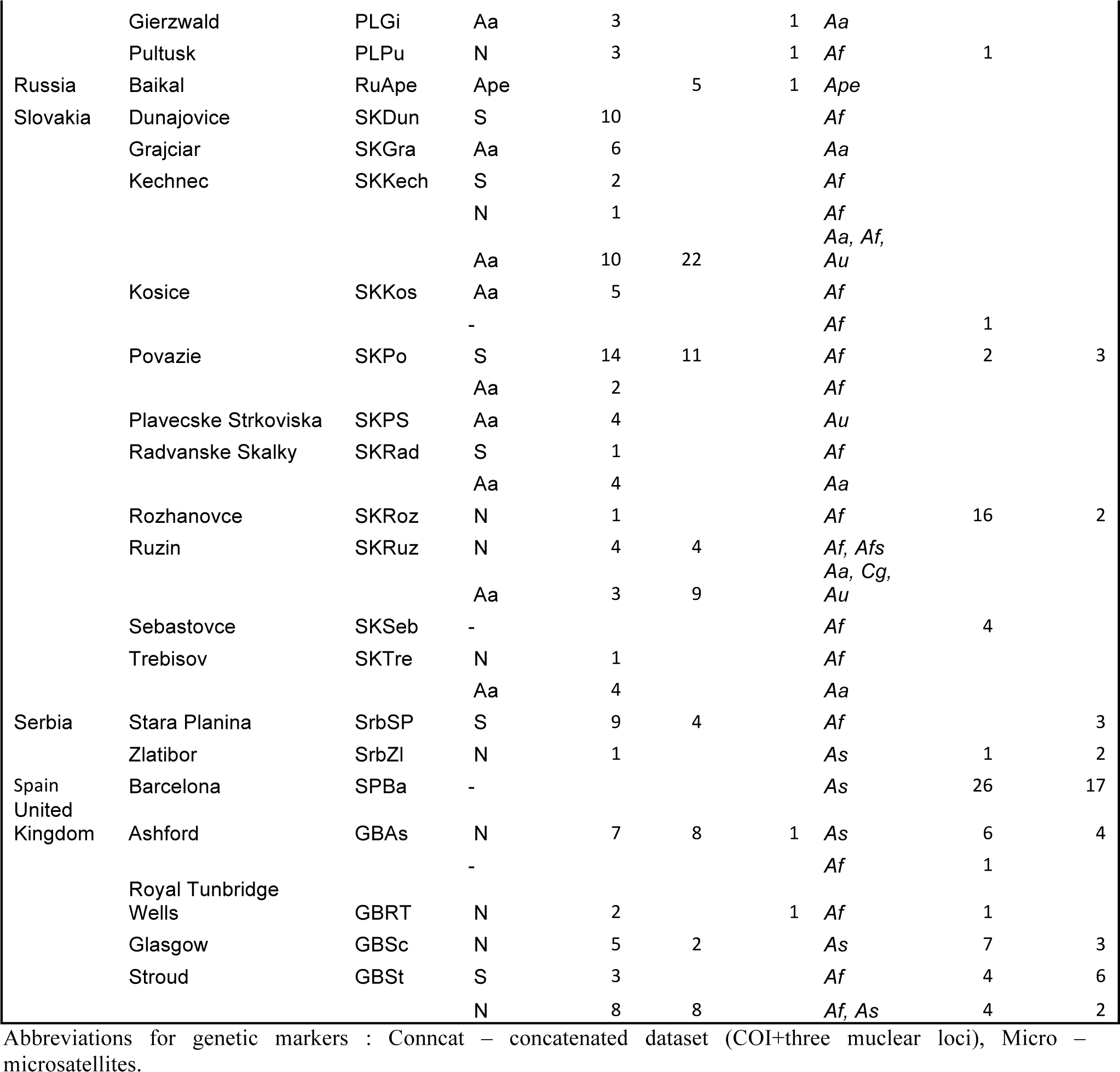

A short (381bp) easily amplifiable COI fragment was used for the basic screening of the total sample set. Phylogenetic analyses clustered the sequences into three well supported lineages (Fig. 2), designated here as N (nonspecific lineage associated with *A. flavicollis* and *sylvaticus*), S (specific lineage only found on *A. flavicollis*) and Aa (a lineage with strong affinity to *A. agrarius* and *uralensis*). The latter lineage was also found on a few *A. flavicollis* and *Clethrionomys glareolus* individuals. The N, S and Aa lineages correspond to the A, B and C clades identified by Štefka & Hypša (2008), respectively. Lice from the S and N lineages occurred sympatrically across large geographic area (Fig. 2). Several subclades were identified within each lineage, usually with sympatric distribution in N and allopatric in S. On the contrary, the less structured Aa lineage was only found in the eastern part of Europe, concurrently with its primary hosts (*A. agrarius* and *uralensis*).

**Fig. 2:**
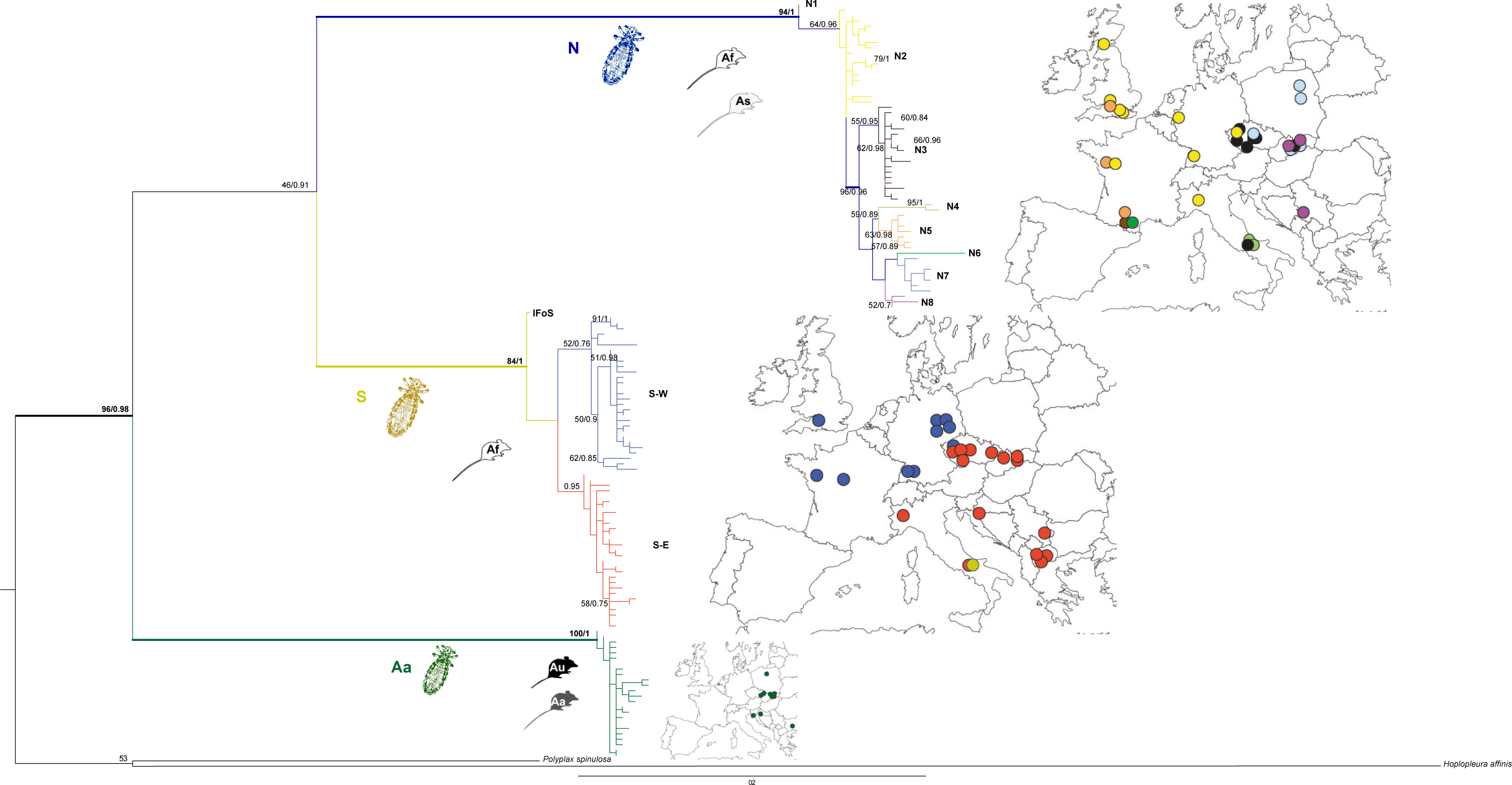
Mitochondrial DNA phylogeny for 556 specimens of *Polyplax serrata*. Maximum Likelihood phylogeny was obtained with PHYML, statistical support (ML bootstrap/Bayesian posterior probability) is provided above clades. Geographical distribution of subclades N and S is provided using matching colours. Abbreviations of clades and host species: N – non-specific clade; S - specific clade; Aa *– Apodemus agrarius* and *uralensis* clade; Af – *A. flavicollis*; As – *A. sylvaticus*; Au – *A. uralensis.*

The relationship between the three main lineages was unclear in this short-sequence matrix, while the analysis of the second matrix consisting of 25 selected samples for which longer COI fragments were concatenated with three nuclear genes, clustered the S and N lineages as sister clades (Fig. 3).

**Fig. 3:**
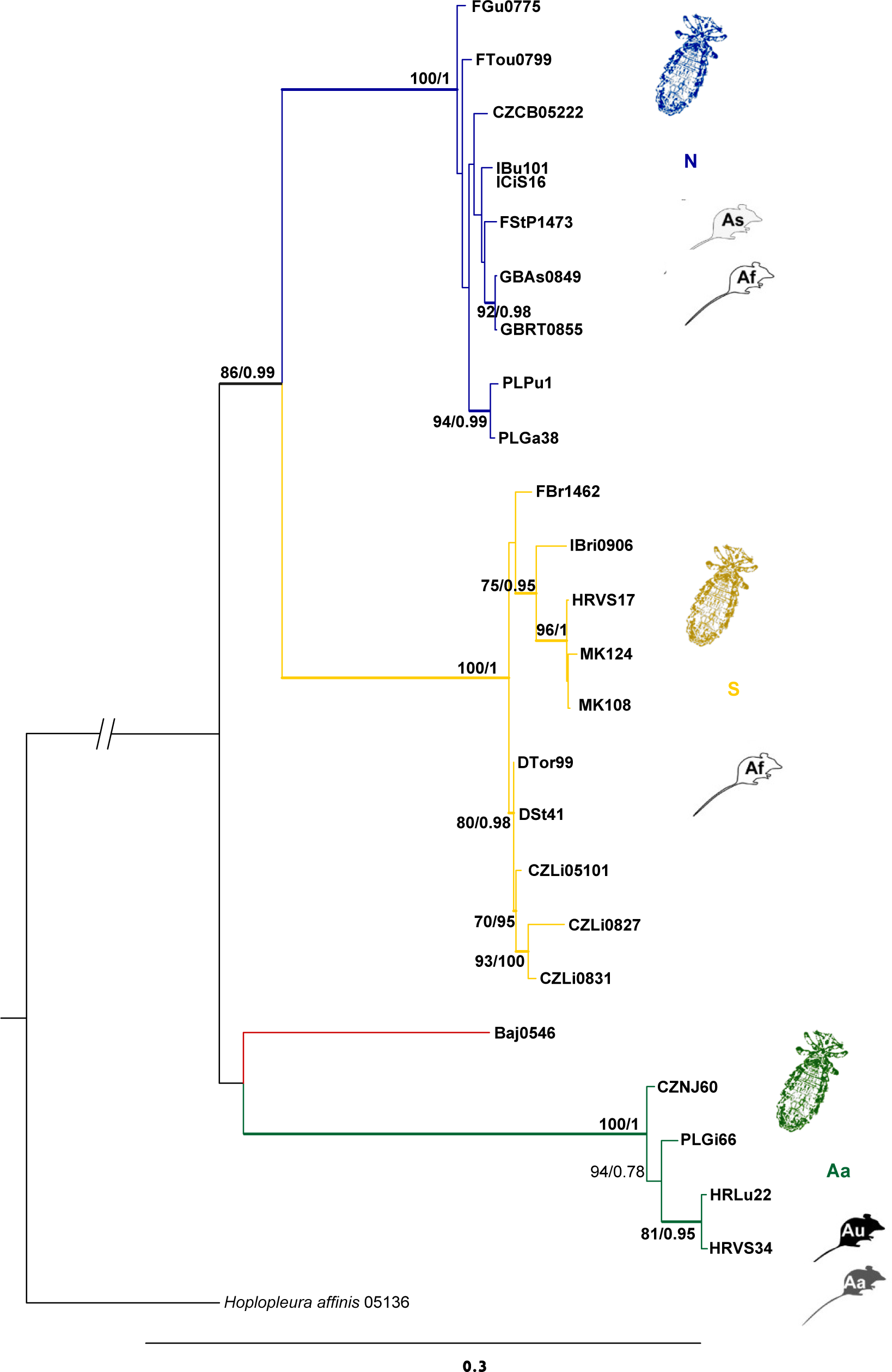
Molecular phylogeny of major *Polyplax serrata* clades basedon a concatenated dataset of four genes (COI, VATP21, hyp and TMEDE6). Maximum Likelihood phylogeny was obtained with PHYML, statistical support (ML bootstrap higher than 50%/Bayesian posterior probability above 0.6) is provided above clades. Abbreviations as in Fig.1.

By collapsing the 556 COI sequences, we retrieved 138 haplotypes that represented from 1 to 97 louse specimens. The overall haplotype network of the main *Polyplax* lineages, corresponded to the phylogenetic topologies (Fig. S2, Supporting information). The S-lineage haplotypes split into two subnetworks (exceeding 95% connection limit) (Fig. 4), representing two geographically distinct populations, hereafter designated as Specific East (S-E) and Specific West (S-W). They only overlapped in a narrow contact zone, represented by two localities in the Czech Republic - CZStr and CZVyk. The Italian IFoS haplotype clustered with S-E in the haplotype network.

**Fig. 4:**
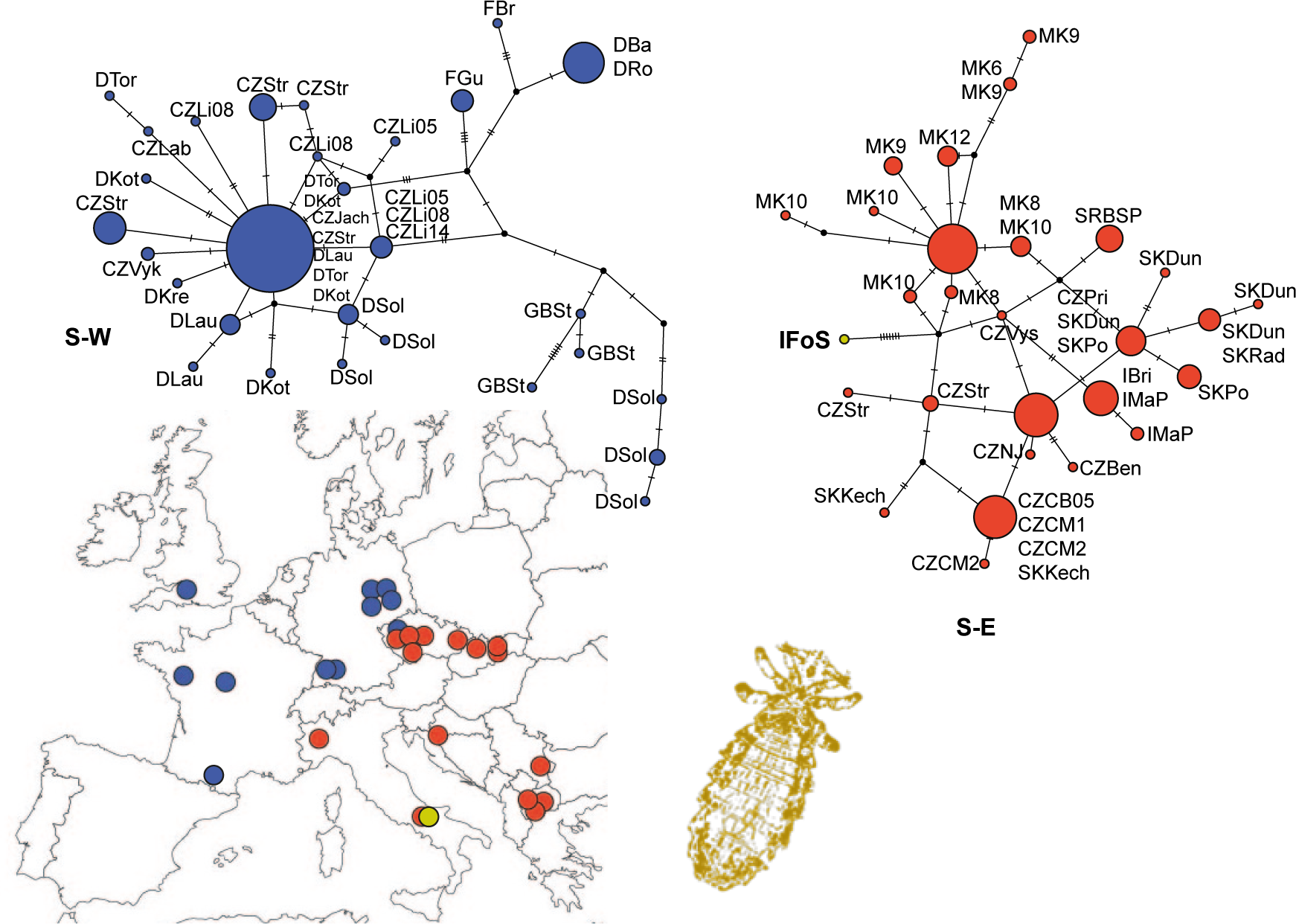
Haplotype networks and distribution map of the S lineage of *Polyplax serrata*. Abbreviations: S-W – Specific West cluster; S-E – Specific East cluster; abbreviations of localities as in Table 1.

Within N lineage, two French haplotypes were isolated from the rest of the network (N1 and N6 in Fig. 5). Unlike the S lineage, no clearcut geographic separation was detected in the N lineage, although certain degree of geographic dependence was discernable. Apart from the widespread cluster ranging from Great Britain to Bulgaria (N2), the N lineage also comprised five clusters with more restricted distribution: Great Britain and France (N5); Italy (N4); Czech Republic (N3); Czech Republic, Slovakia and Poland (N7); and Slovakia and Serbia (N8) (Fig. 5). Majority of the N clusters contained samples from both *A. flavicollis* and *A. sylvaticus*. Out of the three clusters containing multiple haplotypes only the cluster N7 showed narrowed host specificity to *A. flavicollis* (Fig. S3, Supporting information).

**Fig. 5:**
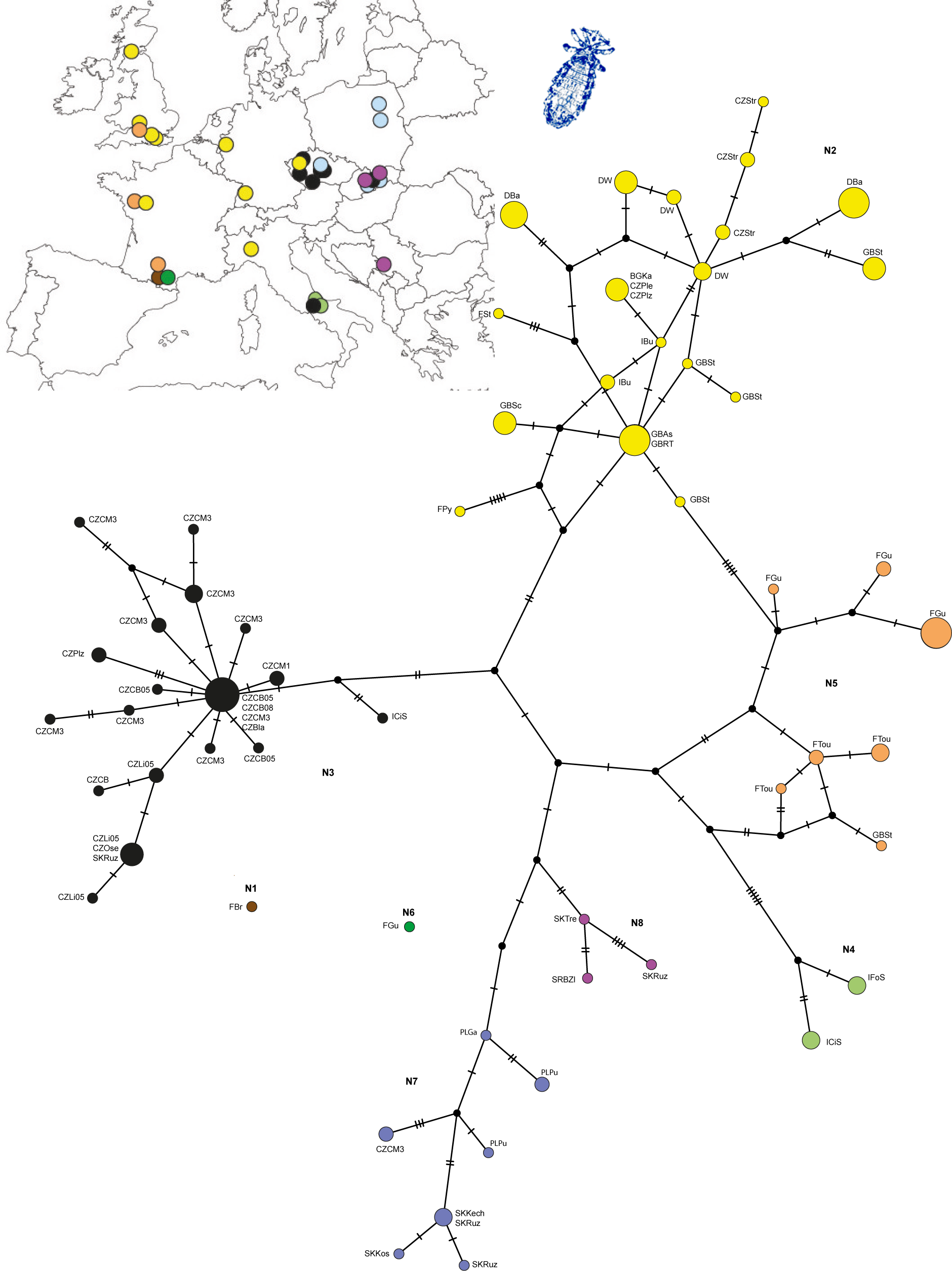
Haplotype network and distribution map of the N lineage of *Polyplax serrata*. Colour code for individual subclades (N1 to N8) as in Fig.1

The network of the Aa lineage was less complex with one central abundant haplotype surrounded by several less common haplotypes, and with tendency of geographically proximate haplotypes to create subclades (Fig. S4, Supporting information).

### Genealogy of *Apodemus sylvaticus* and *A. flavicollis*

For the hosts, D-loop sequences from 230 *A*. *flavicollis* and 93 *A. sylvaticus* mice were obtained and collapsed into 117 and 73 haplotypes, respectively. *A. flavicollis* phylogeny revealed 2 phylogenetically distinct clusters (Af and Bf) largely overlapping in their geographic distribution (Fig. 6) but differing in their abundance.

**Fig. 6:**
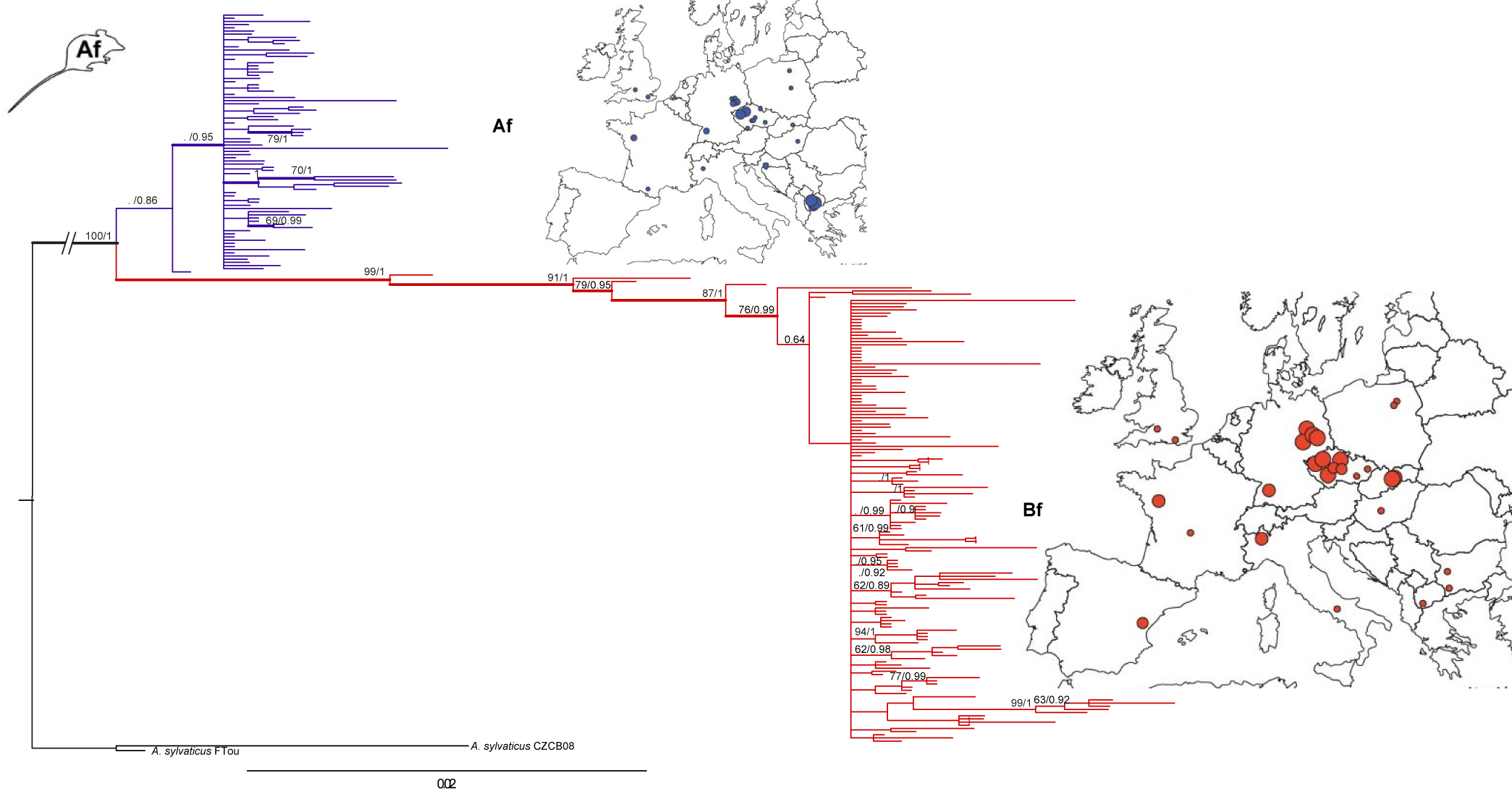
Mitochondrial DNA phylogeny for 230 specimens of *Apodemus flavicollis*. Maximum Likelihood phylogeny was obtained with PHYML, statistical support (ML bootstrap higher than 50%/Bayesian posterior probability above 0.6) is provided above clades. Geographical distribution of subclades Af and Bf is provided using matching colours.

Similar situation was found in *A. sylvaticus*, with phylogenetic tree containing 3 clusters, two of them, As and Bs, with highly sympatric distribution. The less abundant lineage As (n=18) occurred mainly in Great Britain and France, while the more abundant lineage Bs (n=66) extended to Iberian peninsula and was paraphyletic with respect to the Italian-Balkan clade Cc (n=9) (Fig. 7).

**Fig. 7:**
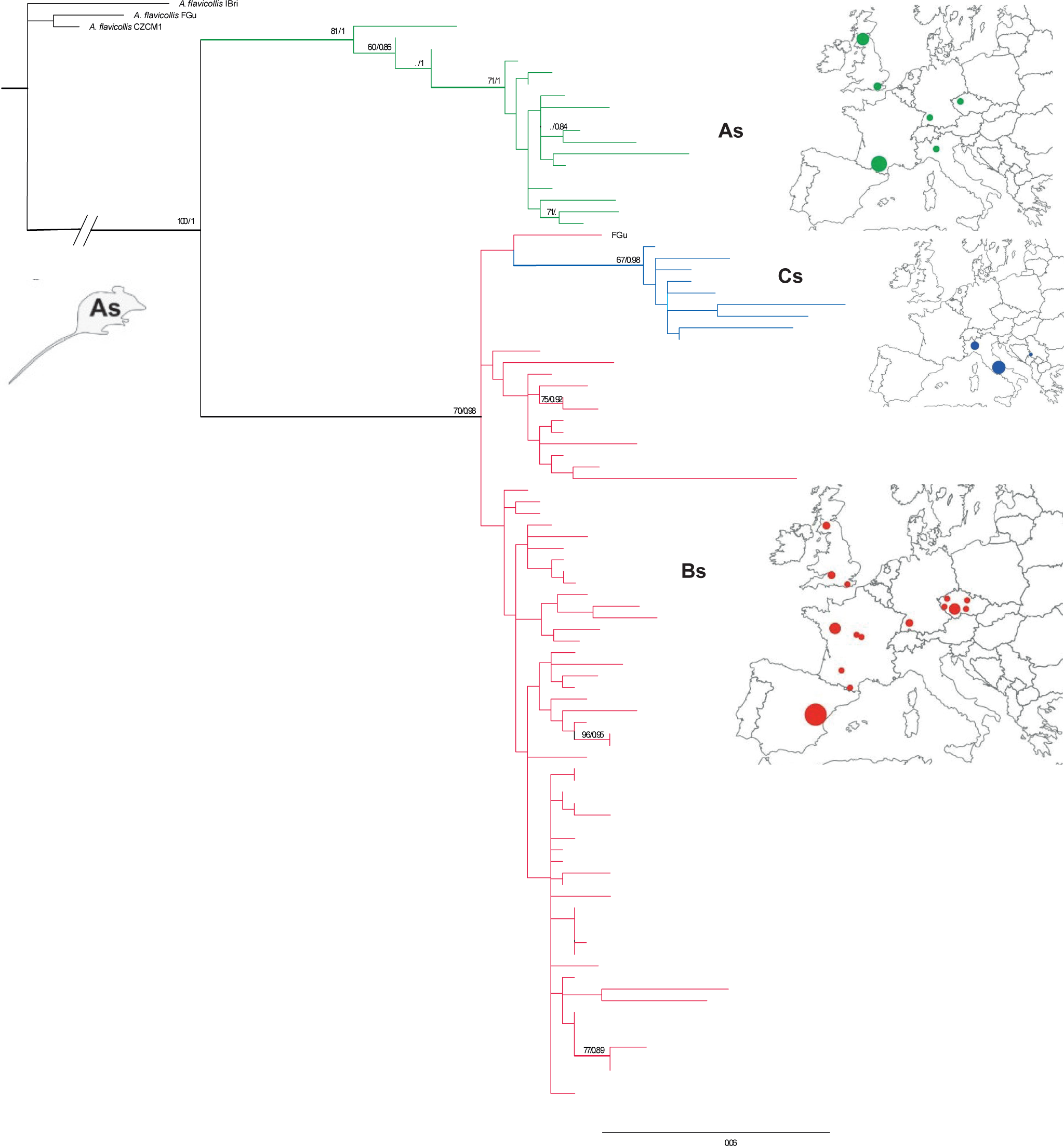
Mitochondrial DNA phylogeny for 93 specimens of *Apodemus sylvaticus*. Maximum Likelihood phylogeny was obtained with PHYML, statistical support (ML bootstrap higher than 50%/Bayesian posterior probability above 0.6) is provided above clades. Geographical distribution of subclades As, Bs and Cs is provided using matching colours.

In both mouse species, the haplotype networks corresponded to the phylogenetic topologies. Genetic distance between the *A. flavicollis* clusters Af and Bf exceeded the 95% connection limit, leading to a split of the samples into two subnetworks. The cluster Bf displayed a complicated inner arrangement, while the cluster Af formed a simpler star-like structure (Fig. S5, Supporting information). Two separated subnetworks, As and Bs+Cs, were also obtained for the *A. sylvaticus* (Fig. S6, Supporting information). Similar to the *A. flavicollis* Af, no major haplotype was present in the less numerous cluster As. The Italian-Balkan clade Cs was separated from the clade Bs by 12 mutation steps, and the clade Bs created complex architecture with network components similar to the clade Bf in *A. flavicollis*.

### mtDNA diversity of *Polyplax* and *Apodemus* populations

#### Polyplax

European louse populations were grouped according to the potential glacial refugia of their hosts, and the areas to which they spread after climatic change (Fig. S1, Supporting information). Samples from the central and western part of Europe (except for the Iberian peninsula, region IV) showed high levels of haplotype and nucleotide diversities for both S and N lineages (Table 2). Tests for recent demographic changes (neutrality indices - Tajima´s D, Fu and Li´s D*, Fu and Li´s F*, Fu´s Fs and R_2_), produced significantly negative values for these regions. Such values were probably caused by an admixture of several parasite lineages after their expansion from distant refugia. In contrast, the lowest levels of the haplotype and nucleotide diversities were detected in the Italian-Balkans (region I) population of the S lineage and for Iberian peninsula (region II) in the N lineage (i.e the areas that according to Michaux *et al.* (2005) served as refugia for *A. flavicollis* and *sylvaticus*).

**Table 2:**
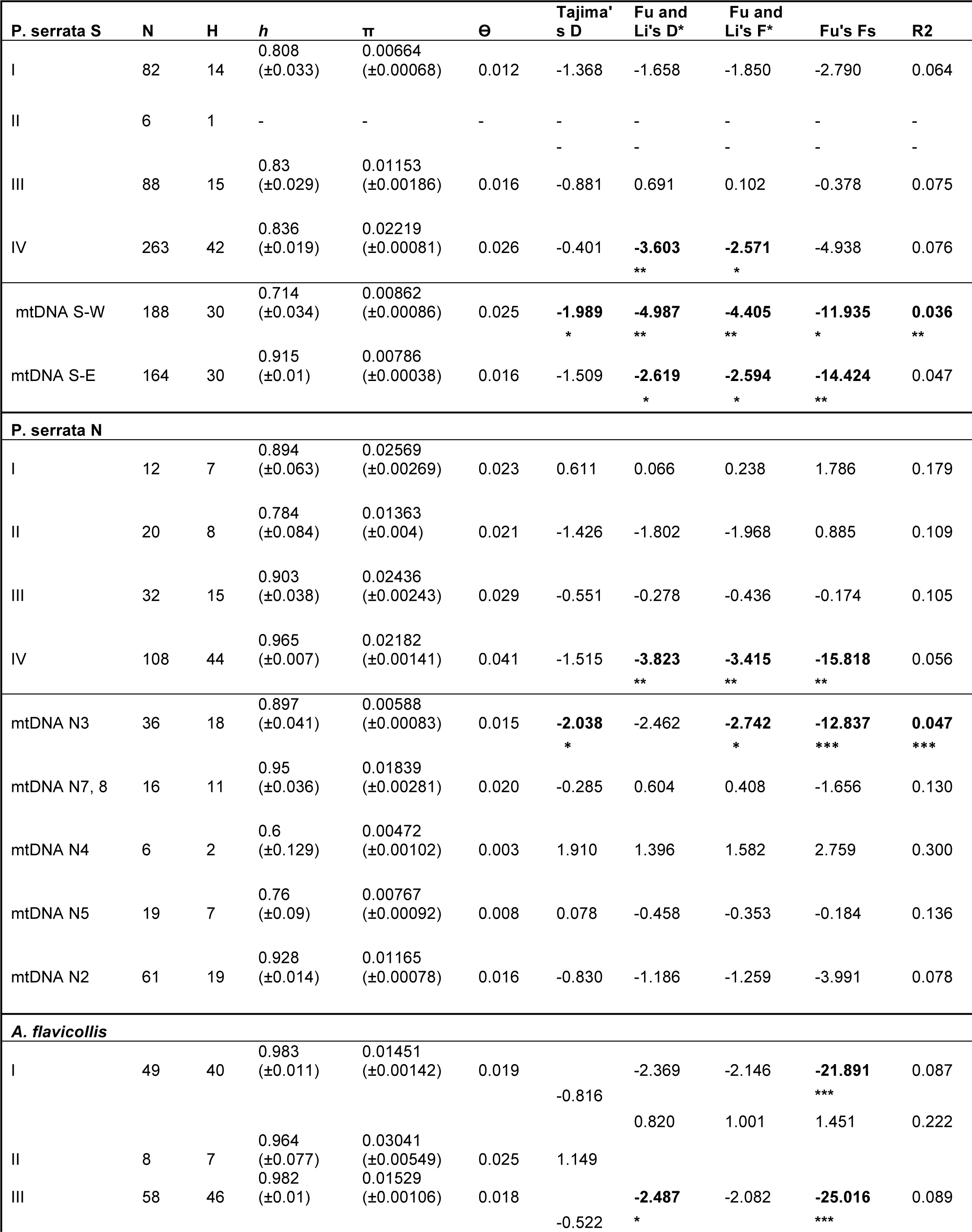
Genetic diversity of populations based on mtDNA.

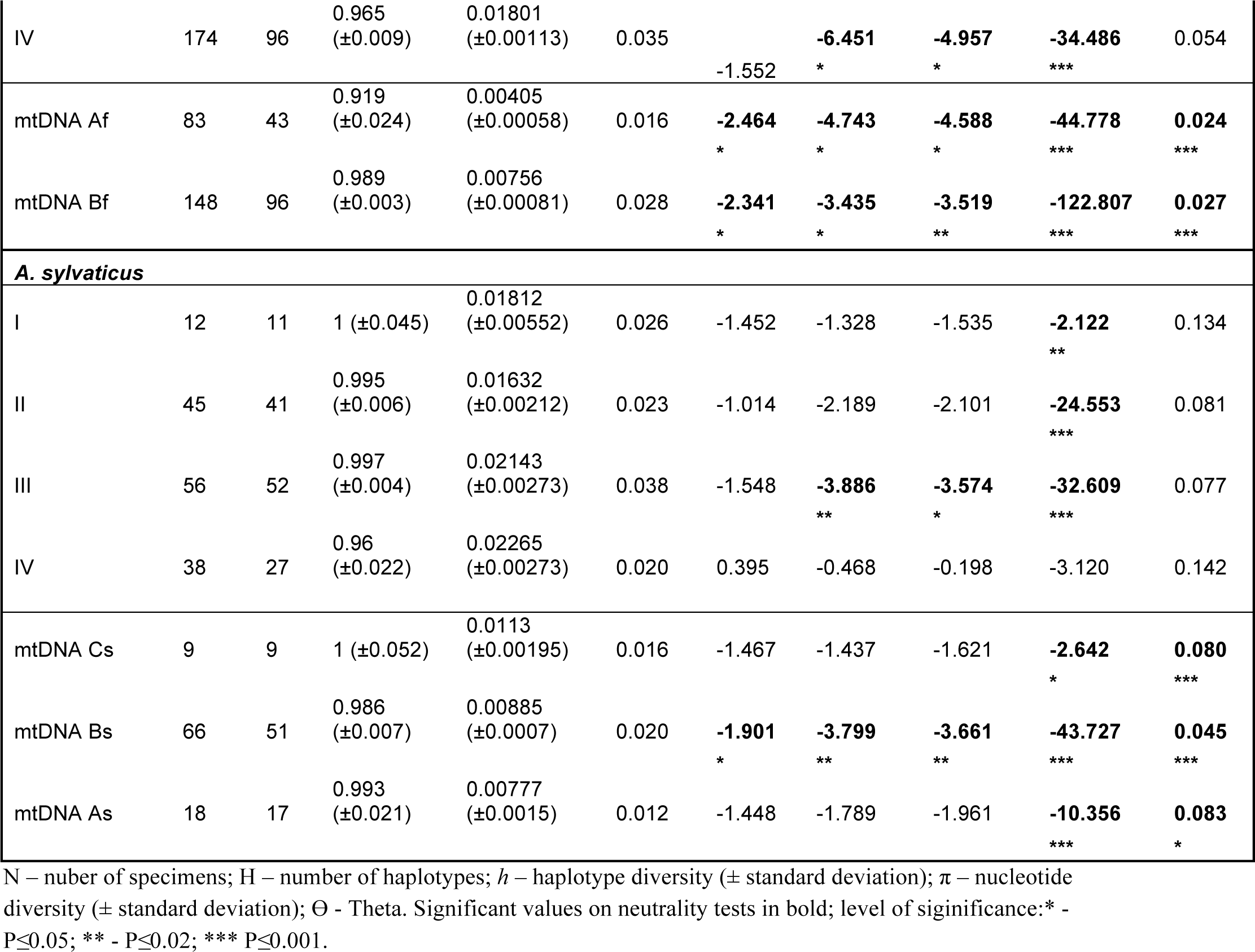

Both S-W and S-E clusters of *Polyplax* showed signatures of potential demographic changes (significantly negative neutrality tests). In the S-W cluster, we detected lower levels of haplotype diversity and higher levels of nucleotide diversity, while higher haplotype and lower nucleotide diversities were seen in the S-E cluster (Table 2). Within the N lineage, cluster from the Italian region (N4) had the lowest nucleotide and haplotype diversities compared to the high values observed in the clusters from Central and Eastern Europe (N7, N8). Deviations from neutral variation (significantly negative values) were only observed in two clusters, one containing the samples from Italy and the Czech Republic (N3) and the other, most numerous, with majority of the samples from Western and Central European localities (N2).

#### Apodemus

Host samples were partitioned into the same four areas as the parasites (Fig. S1, Supporting information). High levels of haplotype diversities were observed in all geographical areas for both *A. flavicollis* and *sylvaticus*. In *A. flavicollis*, samples from the Iberian peninsula (region II) showed higher level of nucleotide diversity and no deviations from neutral processes (non-significant Fu and Li’s tests), while in other regions we obtained statistically significant values of tests (Table 2). In *A. sylvaticus*, populations from Italian and Balkan regions and from the Iberian peninsula showed signs of demographic changes (significant Fu and Li’s test results), but the results were not significant for Central and Western Europe (region IV) (Table 2). When individual clusters within each host species were analysed separately, several of them showed departures from neutrality (namely both *A. flavicollis* mtDNA lineages Af and Bf and lineage Bs of *A. sylvaticus*). Higher values of *h* and π were apparent in the more numerous lineage Bf of *A. flavicollis* and in Cs clade of *A. sylvaticus* (Table 2).

AMOVA analyses of mtDNA datasets organised geographically in the same manner as for the DNASP (Fig. S1, Supporting information) revealed different patterns of population structure for the host and the parasite. Majority of the host variation was distributed within the populations (Table 3), while the *Polyplax* S and N lineages showed significant values among the populations. Significant values were also found among mtDNA clusters of *Polyplax* S-W and S-E lineages. Fixation indexes of both parasites and hosts except of ϕCT of *A. sylvaticus* were significant (Table 3)

**Table 3:**
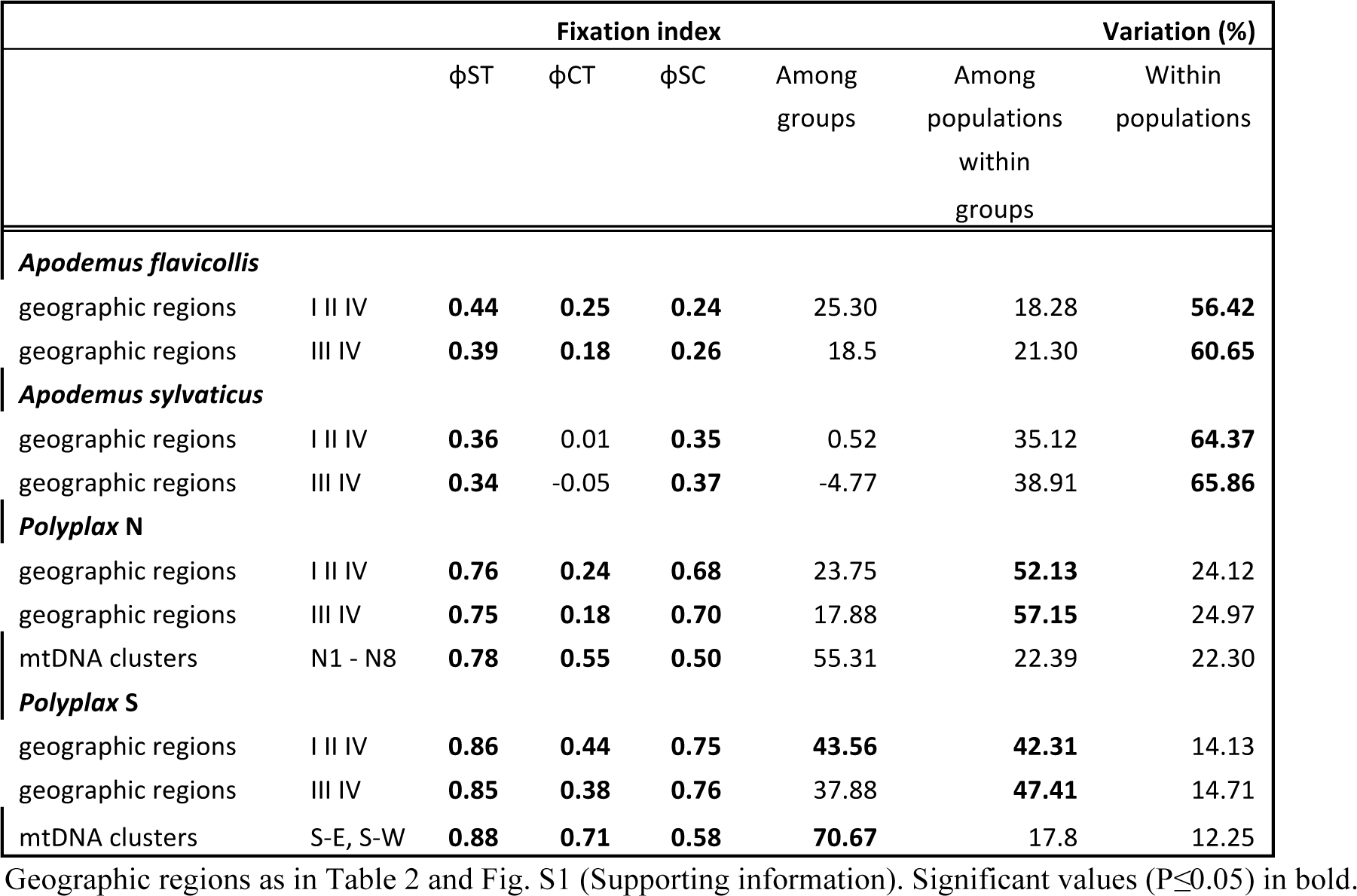
Analysis of Molecular Variance for hierarchically organised poulations of the parasite and two host species.

|  |  | Fixation index |  |  | Variation (%) |  |  |
| --- | --- | --- | --- | --- | --- | --- | --- |
| | | $\phi_{ST}$ | $\phi_{CT}$ | $\phi_{SC}$ | Among groups | Among populations within groups | Within populations |
| ***Apodemus flavicollis*** |  |  |  |  |  |  |  |
| geographic regions | I II IV | **0.44** | **0.25** | **0.24** | 25.30 | 18.28 | **56.42** |
| geographic regions | III IV | **0.39** | **0.18** | **0.26** | 18.5 | 21.30 | **60.65** |
| ***Apodemus sylvaticus*** |  |  |  |  |  |  |  |
| geographic regions | I II IV | **0.36** | 0.01 | **0.35** | 0.52 | 35.12 | **64.37** |
| geographic regions | III IV | **0.34** | -0.05 | **0.37** | -4.77 | 38.91 | **65.86** |
| ***Polyplax N*** |  |  |  |  |  |  |  |
| geographic regions | I II IV | **0.76** | **0.24** | **0.68** | 23.75 | **52.13** | 24.12 |
| geographic regions | III IV | **0.75** | **0.18** | **0.70** | 17.88 | **57.15** | 24.97 |
| mtDNA clusters | N1 - N8 | **0.78** | **0.55** | **0.50** | 55.31 | 22.39 | 22.30 |
| ***Polyplax S*** |  |  |  |  |  |  |  |
| geographic regions | I II IV | **0.86** | **0.44** | **0.75** | **43.56** | **42.31** | 14.13 |
| geographic regions | III IV | **0.85** | **0.38** | **0.76** | 37.88 | **47.41** | 14.71 |
| mtDNA clusters | S-E, S-W | **0.88** | **0.71** | **0.58** | **70.67** | 17.8 | 12.25 |

Geographic regions as in Table 2 and Fig. S1 (Supporting information). Significant values ( $P \leq 0.05$ ) in bold.

### Microsatellite Diversity in *Polyplax*

All microsatellite loci were polymorphic in at least 16 out of 32 populations, with 1 to 11 alleles per locus (Table S1, Supporting information). Significant values of linkage disequilibrium (LD), after sequential Bonferroni correction, were found between 1 to 4 pairs of loci in 3 populations (CZVykS, DBaN, CZStrS). The pairs of loci found to be in LD differed among populations. The average per population heterozygosity values were 0.271 for the observed heterozygosity (*Ho*) and 0.408 for the expected heterozygosity (*He*) (Table 4). All populations showed significant deviations from Hardy–Weinberg equilibrium due to heterozygote deficiencies in at least one locus, but none of the loci was out of HWE across all populations (Table S2, Supporting information). The deviations were more frequent in the S lineage than in the N lineage. Pairwise *F*_*ST*_ values between populations (with n ≥ 5) ranged from 0.042 to 0.572 in the S lineage, 0.108 – 0.281 in the N lineage. *F*_*ST*_ values of the lineages were 0.419 for the S lineage and 0.241 for the N lineage (Table S3, Supporting information). Because plots in DiveRsity, visualizing differences in F_ST_, G_ST_, G'_ST_ and D_JOST_ in relation to polymorphism, indicated possible bias in F _ST_due to high mutation rate of the loci (Figs. S7 and S8, Supporting information), G_ST_ and D_JOST_ were selected to complement the statistics in the analyses of population differentiation (results below).

**Table 4:** Observed and expected heterozygosities for *Polyplax serrata* populations.

| Pop | Ho | He |
| --- | --- | --- |
| CZBenS | 0.131 | 0.417 |
| CZCBN | 0.484 | 0.495 |
| CZCM1N | 0.435 | 0.569 |
| CZCM1S | 0.072 | 0.162 |
| CZDouS | 0.219 | 0.285 |
| CZJachS | 0.200 | 0.256 |
| CZLi05N | 0.348 | 0.383 |
| CZLi05S | 0.229 | 0.481 |
| CZStrN | 0.354 | 0.465 |
| CZStrS | 0.199 | 0.299 |
| CZVyKs | 0.202 | 0.335 |
| DBaN | 0.335 | 0.554 |
| DBaS | 0.353 | 0.420 |
| DKotS | 0.050 | 0.073 |
| DLauS | 0.088 | 0.181 |
| DSolS | 0.161 | 0.274 |
| DTorS | 0.218 | 0.269 |
| FGuN | 0.472 | 0.608 |
| FGuS | 0.110 | 0.348 |
| FTouN | 0.451 | 0.545 |
| GBAsN | 0.343 | 0.459 |
| GBStN | 0.539 | 0.625 |
| HRVSS | 0.328 | 0.363 |
| IBriS | 0.174 | 0.403 |
| MK10S | 0.469 | 0.636 |
| MK12S | 0.000 | 0.455 |
| MK8S | 0.425 | 0.602 |
| MK9S | 0.436 | 0.672 |
| PLPuN | 0.141 | 0.373 |
| SKPoS | 0.136 | 0.174 |
| SKRuzN | 0.422 | 0.547 |
| SrbSPS | 0.141 | 0.324 |
| **Average** | **0.271** | **0.408** |

Population abbreviations as in Table 1.

### Microsatellite Diversity in *Apodemus*

Number of alleles in *A. flavicollis* and *sylvaticus* varied from 1-15 alleles per locus with an average of 4 alleles per locus and population (Table S4, Supporting information). After Bonferroni correction none of the loci in the two species showed LD. Average values of *Ho* and *He* per populations were 0.657 and 0.709 for *A. flavicollis* and 0.647 and 0.715 for *A. sylvaticus*, respectively (Table 5). HWE were calculated for populations with n≥5 individuals. In *A. flavicollis*, for which 12 loci were analysed, two populations were in HWE, the rest showed deviations from HWE in 1 to 4 loci, and the German population DLau had 6 loci out of HWE (Table S5, Supporting information). In *A. sylvaticus*, with 17 loci analysed, the British population GBA showed no deviations from HWE, majority of the other populations had 1 to 4 loci out of HWE, the French population FTou had 5 loci and the Spanish population EBa had 11 loci out of HWE (Table S6, Supporting information). Pairwise values ranged from 0.156 to 0.306 in *A. flavicollis* and 0.019 to 0.136 in *A. sylvaticus* (Table S3, Supporting information).

**Table 5:** Observed and expected heterozygosities in *Apodemus flavicollis* and *sylvaticus* populations.

| *Apodemus flavicollis* |  |  | *Apodemus sylvaticus* |  |  |
| --- | --- | --- | --- | --- | --- |
| Pop | Ho | He | Pop | Ho | He |
| **CZBen** | 0.667 | 0.552 | **FTou** | 0.638 | 0.793 |
| **CZCM1** | 0.563 | 0.622 | **FGu** | 0.574 | 0.631 |
| **CZDou** | 0.583 | 0.709 | **GBAs** | 0.750 | 0.664 |
| **Fin** | 0.542 | 0.503 | **GBSt** | 0.594 | 0.663 |
| **Fgu** | 0.639 | 0.767 | **DBa** | 0.718 | 0.729 |
| **lbri** | 0.668 | 0.748 | **Eba** | 0.687 | 0.848 |
| **CZLi08** | 0.638 | 0.818 | **CZPI** | 0.571 | 0.679 |
| **Mk10** | 0.764 | 0.799 | **Total** | 0.647 | 0.715 |
| **MK9** | 0.732 | 0.809 |  |  |  |
| **DBa** | 0.600 | 0.738 |  |  |  |
| **DKrei** | 0.741 | 0.741 |  |  |  |
| **DLau** | 0.660 | 0.752 |  |  |  |
| **DPin** | 0.604 | 0.641 |  |  |  |
| **DSol** | 0.735 | 0.722 |  |  |  |
| **DTor** | 0.740 | 0.734 |  |  |  |
| **Draj** | 0.625 | 0.630 |  |  |  |
| **CZStr** | 0.670 | 0.763 |  |  |  |
| **Total** | 0.657 | 0.709 |  |  |  |

Populations with less than 5 individuals were not listed. Population abbreviations as in Table 1.

### Genetic structure based on nuclear microsatellites

#### Polyplax

PCoA analysis of the whole dataset revealed substantial genetic variation (explained variance: axis 1 - 24.47%, axis 2 – 17.39%, axis 3 – 8.91%) and divided the populations into clusters according to the main mtDNA lineages (N lineage - blue circle, S – yellow circle, Aa – green, Bai – red, Fig. 8). The only discrepancy in the lineage assignment compared to the mtDNA data was represented by a Czech population from Litvínov (CZLi05N - marked in red in Fig.8), which belongs to the N lineage according to the mtDNA data, but clusters together with S populations in the microsatellite analysis.

**Fig. 8:**
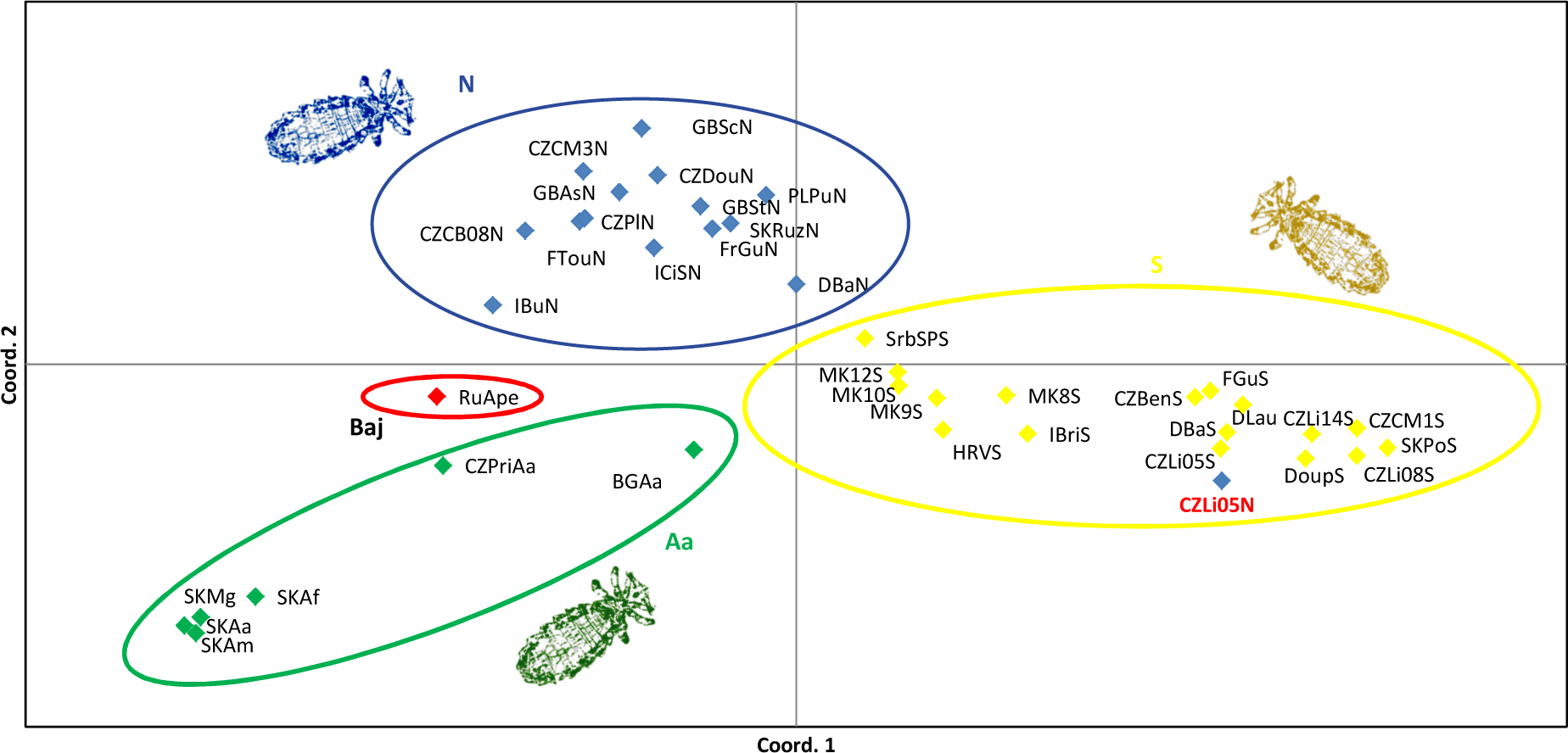
Principal Coordinate Analysis (PCoA) of *Polyplax serrata* populations using microsatellite data. Colours match major lineages used in Fig. 3. Population sample containing mtDNA introgressed from the N lineage (CZLi05N) is highlighted in red. Population abbreviations as in Table 1.

STRUCTURE analysis run for K=2 and 4 yielded highest values of H' in CLUMPP. While results for 2 clusters did not correlate neither with the mtDNA genealogy nor with geographic distribution of the samples, assignment into 4 clusters reflected main mtDNA lineages in the same way as was seen in PCoA (Fig. S9, Supporting information).

BAPS, under the assumption of 4 or 5 clusters (K=4, 5), showed slightly different pattern from Structure and PCoA (Fig. S9, Supporting information). One cluster comprised populations from the N lineage together with Baikal population from *A. peninsulae* (RuApe), two (K=4) or three (K=5) clusters involved populations from the S lineage together with CZLi05N population, Aa populations clustered separately into the last cluster.

On the intra-lineage level, PCoA analysis of individuals belonging to S (axe 1 – 17.39%, axe 2 – 15.62%, axe 3 – 8.91%) and N (axis 1 – 10.26%, axis 2 – 7.8%, axis 3 – 5.86%) lineages showed that in majority of the cases lice sampled from the same geographic locality formed compact structures, and populations located geographically close to each other often showed genetic proximity (Fig. 9a, b). The trend was more pronounced in the S lineage compared to N. PCoA analysis run on a population level (S lineage - axis 1 – 27%, axis 2 – 13.3%, axis 3 – 11.51%; N lineage - axis 1 – 17.66%, axis 2 – 14.33%, axis 3 – 12.76%) revealed further differences between the S and N lineages (Fig. 9c, d). S lineage populations created clusters according to their geographic origin (e.g. Italo-Balkan, central Europe, North-East Germany), whereas among the N lineage populations fractional geographic clustering was discernible, but it did not create such explicit units like in the S lineage.

**Fig. 9:**
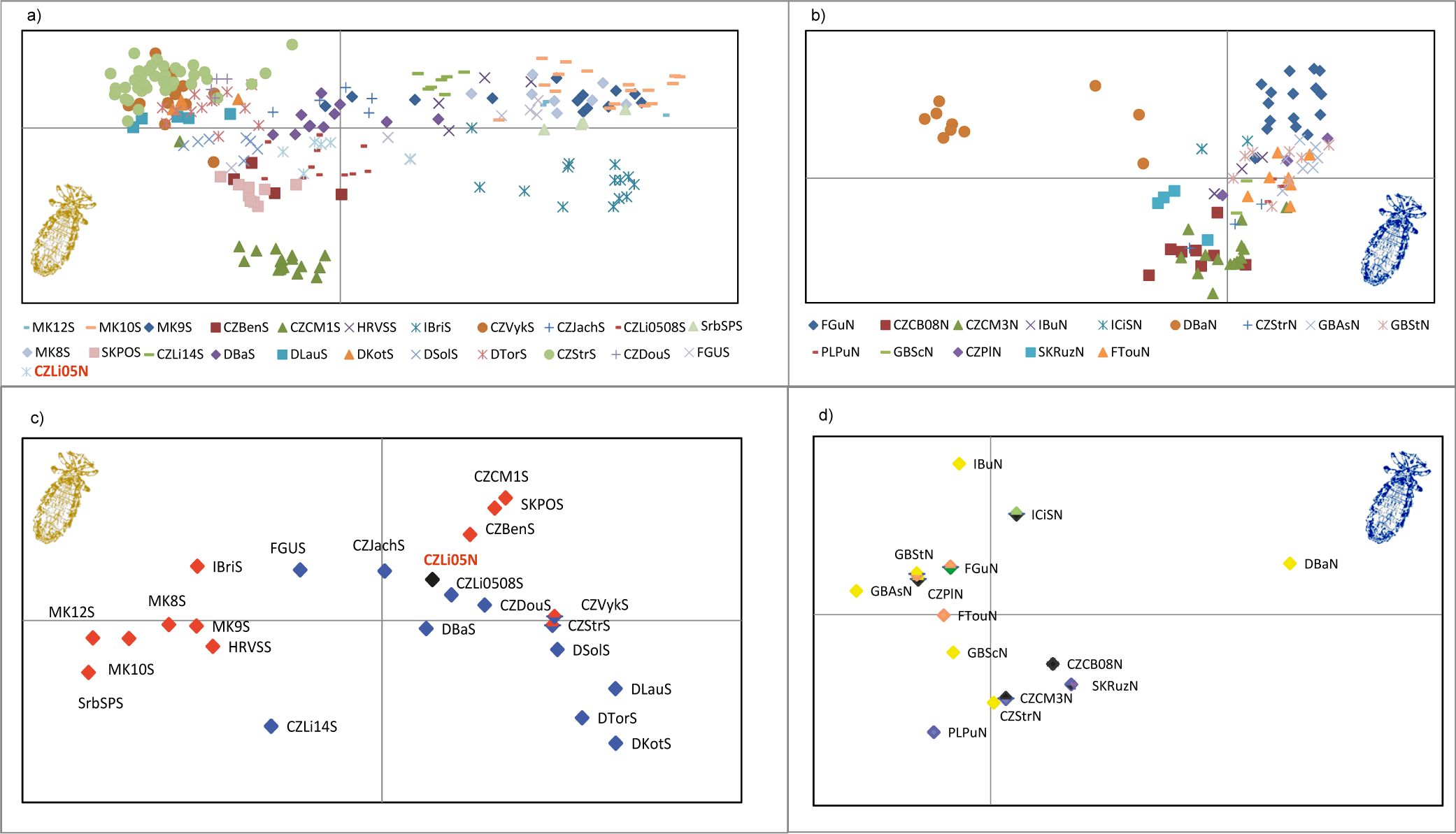
PCoA of *Polyplax serrata* individuals a) and b) and poulations c) and d) belonging to S and N clades (respectively) using microsatellite data. Colours in c) and d) match major lineages used in Fig. 2. Specimens containing mtDNA introgressed from the N lineage (CZLi05N) are highlighted in red. Population abbreviations as in Table 1.

Genetic differentiation between the S and N lineages was also obvious from the NJ tree built using Nei´s DA distances (Fig. 10) and from the assignment test performed in GenAlEx (results not shown). Both analyses also confirmed the microsatellite clustering of CZLi05N to S instead of N lineage. Similarly to PCoA, NJ tree often provided statistical support for clustering between proximate populations, but generally lacked support on a higher geographical level. Similar picture was provided by Bayesian clustering methods (Fig. S10, Supporting information) indicating that intra-lineage hierarchical structure remained unclear with microsatellite data.

**Fig. 10:**
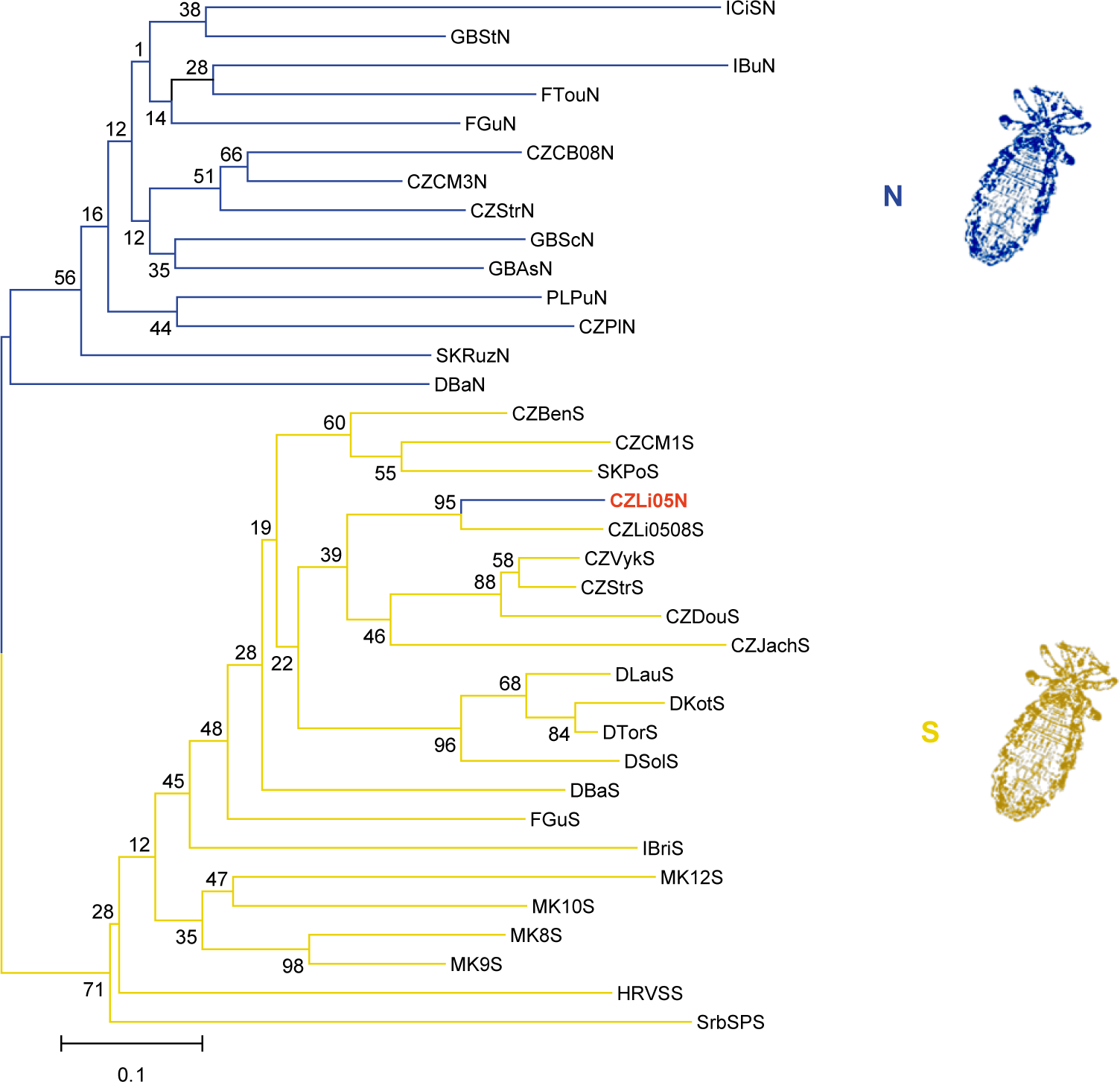
Neighbor-joining analysis of populations of *Polyplax serrata* S and N clades using microsatellite data. Colours match Figs. 2 and 3. Population sample containing mtDNA introgressed from the N lineage (CZLi05N) is highlighted in red. Population abbreviations as in Table 1.

#### Apodemus

Several types of analyses (Structure, PCoA, BAPS, TREEVIEW) performed on a set of 7 microsatellite loci shared by both host species confirmed in concordance with mtDNA results that *A. flavicollis* and *A. sylvaticus* represent two separated species units (results not shown). On the intra-specific level, PCoA analyses performed on separated datasets of *A. flavicollis* (axis 1 – 8.17%, 2 – 3.31%, 3 – 3.04%) and *A. sylvaticus* (axe 1 – 5.47%, 2 – 4.73%, 3 – 4.62%) showed that host individuals belonging to different mtDNA subclades did not form discernible clusters when retrieved from sympatric localities. Geographic populations (localities) were more admixed than in the parasites and did not cluster together (Fig. 11a, b). Similar results were obtained also in Structure and BAPS (Fig. S10, Supporting information) PCoA performed on a population level (Af – axis 1 – 25.83%, 2 – 9.51%, 3 – 8.38%; As – axis 1 – 20.13%, 2 – 17.5%, 3 – 12.02%) (Fig. 11c, d), and PopTreeW analysis (Fig. S11, Supporting information), showed emerging formation of several genetic lineages, which however did not reflect mtDNA genealogy and showed only limited correspondence with geography (e.g. GB and FR populations in *A. sylvaticus*, Fig. 11d).

**Fig. 11:**
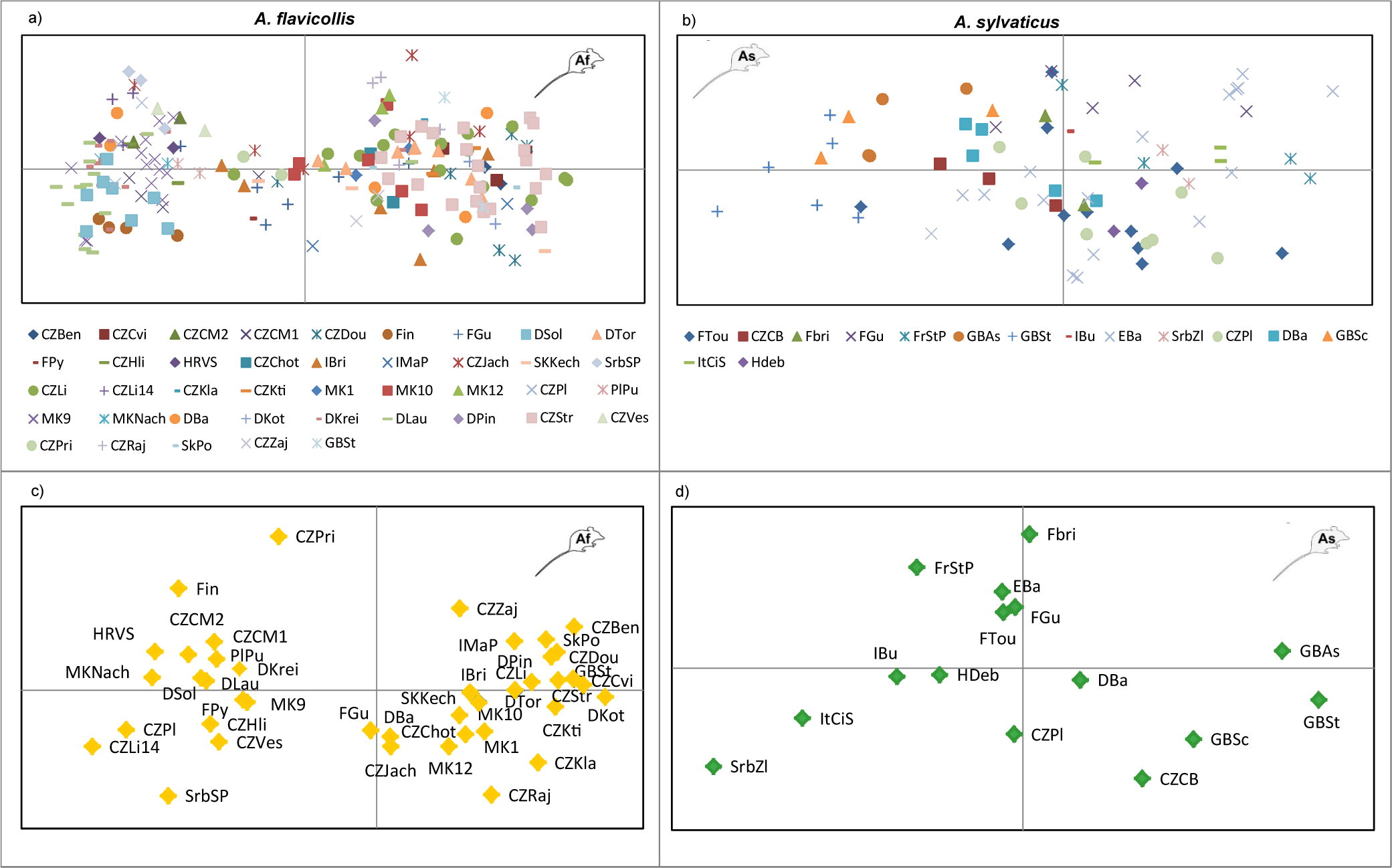
PCoA of *Apodemus flavicollis* a) and *A. sylvaticus* b) individuals and poulations c) and d) (respectively) using microsatellite data. Population abbreviations as in Table 1.

#### Spatial structure of parasites and hosts

The Mantel tests correlating genetic pairwise matrices (*F*_*ST*_, *G*_*ST*_ and *D*_*JOST*_)with geographic distances revealed differences in organisation of population structure between the hosts and the two parasite lineages. But they also produced different results depending on the statistics used. F_ST_ tests found significant IBD only within *A. sylvaticus* (Fig. S12, Supporting information). G_ST_ tests were statistically significant for *Polyplax* S lineage and for *A. sylvaticus*, whereas D_JOST_ test was significant only for *Polyplax* S lineage (Fig.12). Isolation-by-distance analyses measuring the correlation between Euclidean distances (performed on the level of individuals) and geographic distances were significant for both S and N lineages, with a larger correlation coefficient for the S lineage (Fig. S13, Supporting information).

**Fig. 12:**
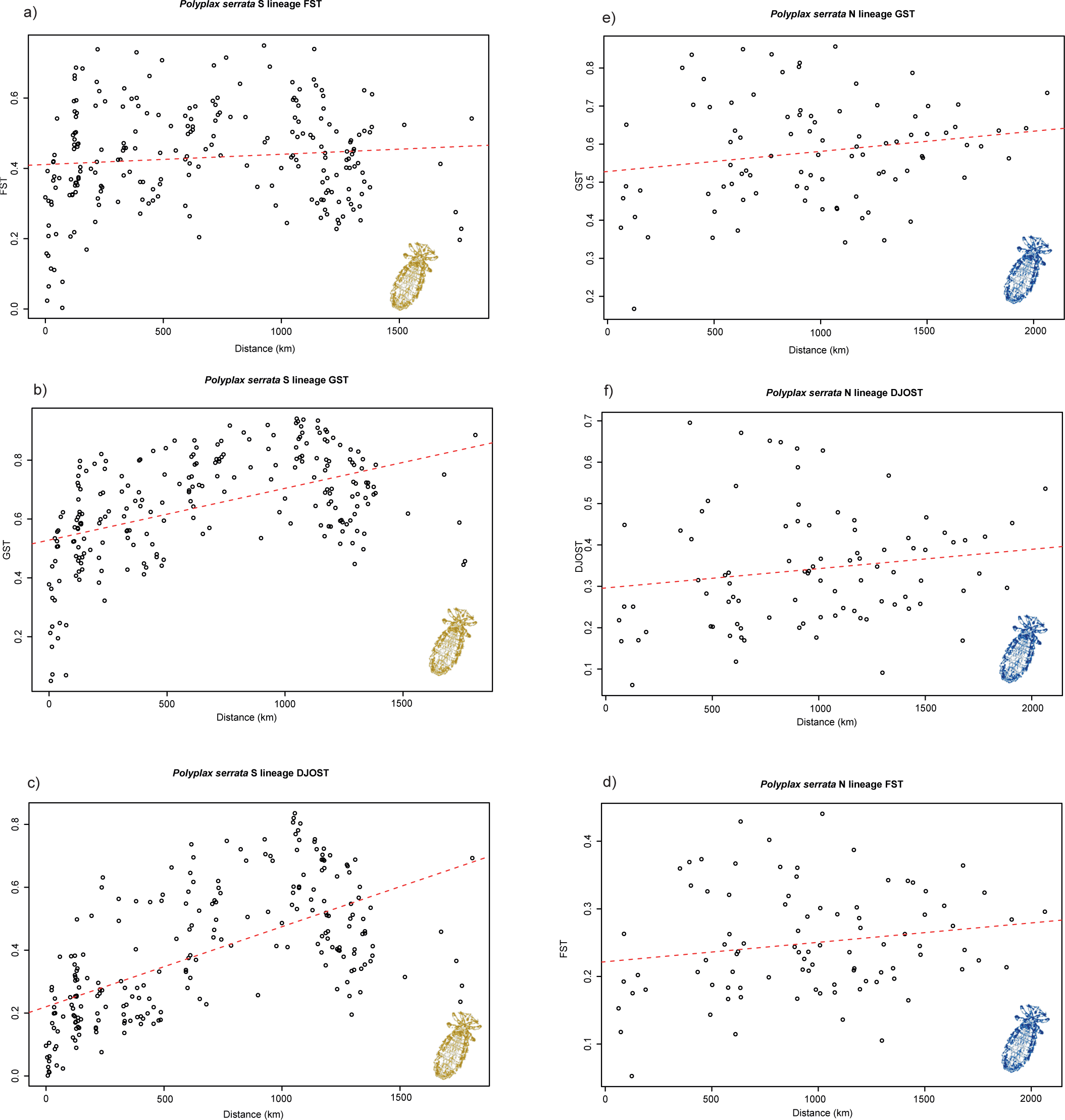
Isolation by distance in the populations of *Polyplax serrata* S (a to c, in yellow) and N lineages (d to e, in blue). Mantel test for correlation between pairwise F_st_ (a and d), G_ST_ (b and d) and D_JOST_ (c and f) indices and geographical distances is marked by red dashed line. Corelation is significant (P<0.005) for d) G _ST_ and c) DJOST of *Polyplax serrata* S.

Autocorrelation coefficient (*r*), used to evaluate the effect of IBD on different geographic scales, revealed positive significant autocorrelation in all evaluated organisms (*Polyplax* lineages S and N, *A. sylvaticus* and *A. flavicollis*) (Fig. 13). However, the spatial extent and the strength of autocorrelation differed between organisms. In lice *r* became non-significant at 350 and 400km for the N and S lineages, respectively, whereas in the hosts at 400 km for *A. flavicollis* and at 550 km for *A. sylvaticus*. More importantly, *r* dropped down to significantly negative values with increasing geographic distance of 600-1200 km in both *Polyplax* lineages, whereas in hosts it remained nonsignificant for majority of the range between 500 and 1200 km. Positive autocorrelation coefficient was 10 times lower at the shortest distance range in *A. flavicollis* than in *A. sylvaticus*, which corresponded with nonsignificant results of Mantel tests in *A. flavicollis*. On the contrary, the highest values of *r* in *Polyplax* lineages were two times greater than that of *A. sylvaticus.*

**Fig. 13:**
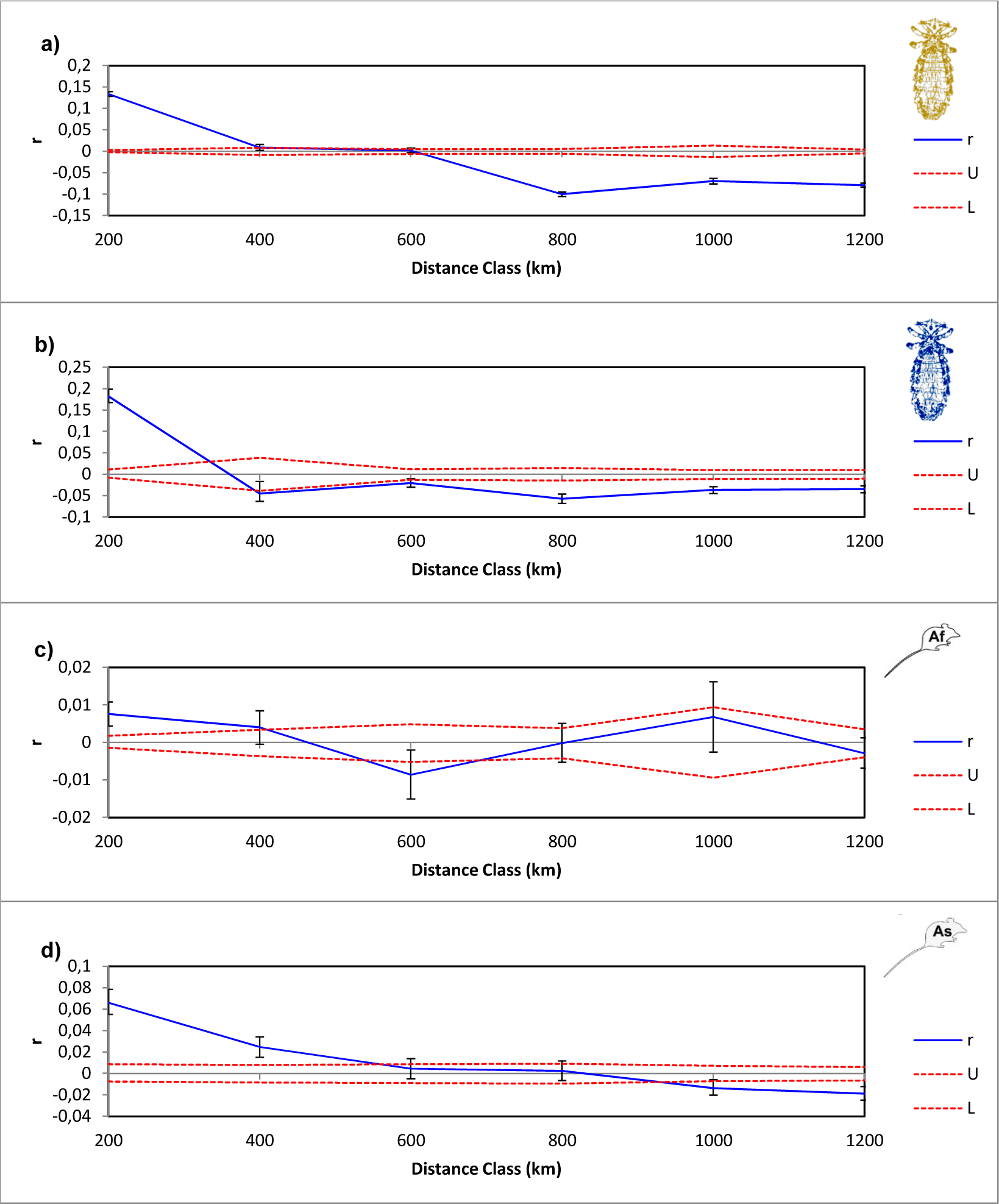
Plots of genetic autocorrelation coefficient (r) across increasing geographic distance class sizes for *Polyplax serrata* S lineage a), *Polyplax serrata* N lineage b), Apodemus flavicollis c) and *A. sylvaticus* d). Error bars indicate the 95% confidence interval around the observed r values, and red dashed lines mark the upper (U) and lower (L) 95% confidence interval for the null hypothesis of no spatial autocorrelation (*r* = 0).

### Differences in population diversities between S and N lineages of *Polyplax*

Microsatellite data that reflected very well geographical distribution of genetic structure in the parasite were used to verify Nadler´s rule using populations of the S and N lineages as representatives of the specialist and generalist parasitic strategies. According to our prediction, *F*_*ST*_ and *H* indices calculated for each of the two lineages revealed lower genetic diversity and stronger population structure of the S lineage. F_ST_ index was statistically lower for N lineage (0.241) than for S lineage (0.46) (15 000 permutations). Conversely, the *H* index was markedly higher for populations of the N lineage (0.587) than for S populations (0.389) (15 000 permutations). More detailed study of both lineages performed on 7 pairs of sympatric (or closely located populations) (Fig. 14) showed in all pair-wise comparisons higher values of *H* for N populations than for S.

**Fig. 14:**
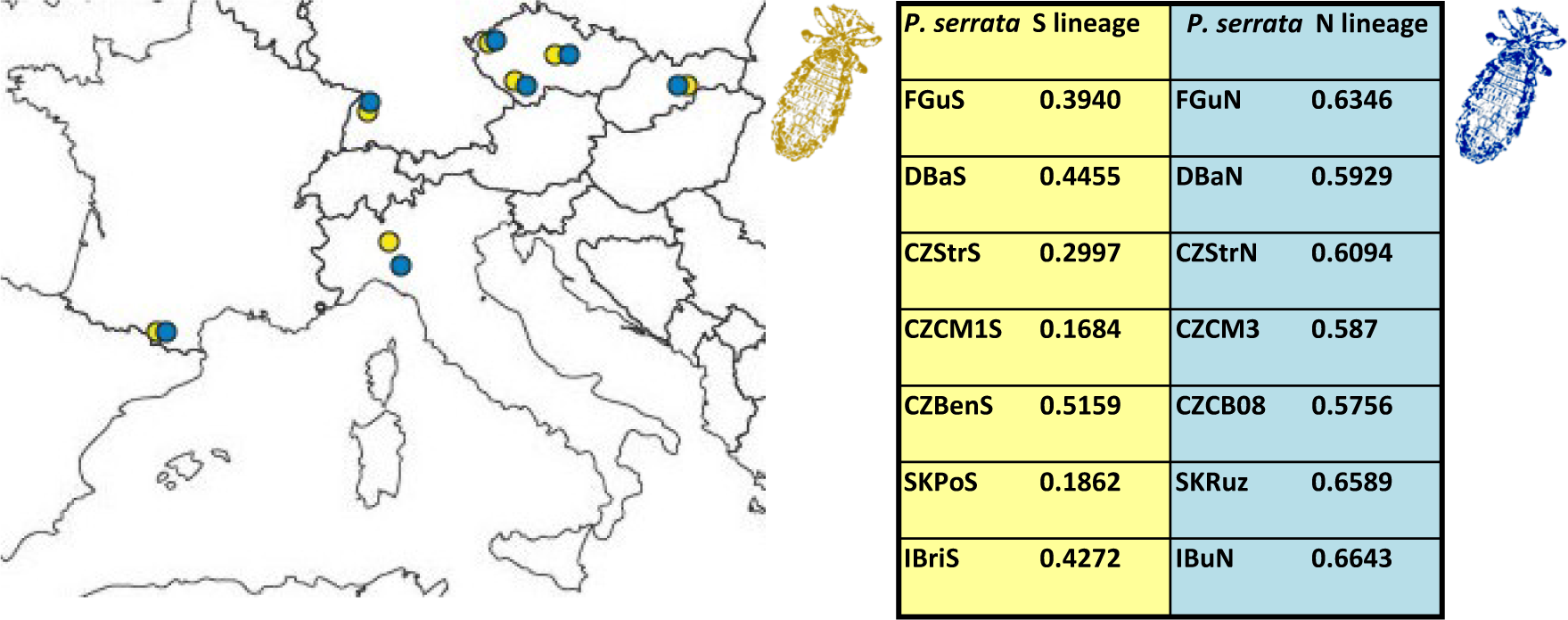
Gene diversity (*H*) and geographic distribution for 7 pairs of sympatric S and N lineage populations. Colour code as in Fig. 2. Population abbreviations as in Table 1.

## Discussion

On the *Apodemus*/*Polyplax* model, we demonstrate that co-evolutionary processes, when viewed from a broad-scale population perspective, may produce surprisingly complex and intriguing patterns. At an overall generalized level, they conform to the traditionally held views that parasites’ phylogenies and genealogies are strongly determined by their hosts, and that populations of parasites have lower genetic connectivity and are more structured than those of the hosts. However, at a more subtle level, the structure, genetic diversity and host specificity of the parasite’s populations strikingly differ even between closely related sister clades. We document this by comparison between the specific lineage S, with low genetic diversity and higher level of isolation by distance, and the more generalist N lineage found on two host species. However, the most striking instance is provided by the sharp difference in the postglacial colonization process between the *A. flavicollis* and its specific parasite, the S lineage of *Polyplax*. For the host, the encounter of populations from different refugia resulted in a largely panmictic European population. In contrast, the louse populations remained genetically separated, with only a narrow contact zone (discussed below). The complexity of the whole system is further increased by various unique genetic events, such as a mitochondrial introgression of the N louse clade into a single population of the other clade.

### *General phylogenetic and co-evolutionary pattern in* Apodemus/Polyplax *system*

The distribution of clades and haplotypes within the two *Apodemus* species reflects biogeographic and climatic changes during Quaternary glaciation. In *A. flavicollis* (Fig. 6), the clade Bf likely originated in the Russian-Ukrainian refugium or the Balkan region, as was suggested by Michaux *et al.* (2005), and then expanded westwards. Clade Af possibly persisted in the Balkan region, where it survived till now, but also expanded to the north. This scenario is also consistent with the neutrality tests. Significant values of Fu’s Fs test for Balkan region and deviations from neutrality in Central, Western and Northern Europe (region IV, Table 2) are most likely caused by simultaneous occurrence of two different mtDNA lineages. For *A. sylvaticus*, Iberian peninsula or Southern France was the main refugium, from where clades As and Bs recolonized Europe, and the Italian-Balkan region served as a possible refugium for the geographically restricted subclade Cs (Fig. 7). This scenario is in agreement with the significant Fu’s Fs, especially in the Italian region (Tab.2). The postglacial histories inferred for both hosts also correspond to the results of Michaux *et al.* (2003, 2005) based on Cytochrome b (*cytB*) sequences. In each of the host species, we found simultaneous distribution of mtDNA clades on the majority of European sites, which suggests post-glacial admixture of populations spreading from the different refugia, as further supported by AMOVA and microsatellite results. PCoA and Structure analyses (Fig. 11; Fig S10, Supporting information) confirmed that in most cases different mtDNA clusters of the same species when living in sympatry now form panmictic populations.

For the lice, the basic division into three major clusters (S, N and Aa, Fig. 2) corresponded to the topology suggested by Štefka & Hypša (2008). Here, we retrieved the S and N lineages as monophyletic sister clades (Fig. 3), clearly separated by the microsatellite data (Fig. 8). The two lineages are likely cryptic species with concurrent geographical distribution, and thus provide an excellent background for a detailed comparative analysis from the population genetic perspective. The third European lineage, Aa, probably represents a recent colonizer from eastern Palearctic, which is documented by the shape of its haplotype network (Fig. S4, Supporting information), close relationship to Ape lineage from the Baikal Lake, and by known history of its *A. agrarius* and *uralensis* hosts (Suzuki *et al.* 2008).

Haplotype diversity and neutrality tests for S and N clades showing similar patterns to the hosts (Table 2), indicate that the lice retreated to refugia and recolonized the northern areas together with their hosts. However, despite this, *P. serrata* and *Apodemus* mice show only limited degree of concordance in phylogeographic distribution of their respective clades (see Figs. 2, 6 and 7, and the discussion below). Limited concordance was reported also by du Toit *et al.* (2013) between *Polyplax arvicanthis* and *Rhabdomys* mice. However, *P. arvicanthis* has two times higher abundance and five times higher prevalence than *P. serrata*, and thus reaches higher effective population sizes and its genealogy is less diversified than the genealogy of its host. Moreover, the four *Rhabdomys* species have a parapatric distribution with narrow contact zones instead of the sympatric occurrence found for the populations of *A. sylvaticus* and *A. flavicollis.* The *Rhabdomys*/*P. arvicanthis* thus provides a contrasting rather than comparative system to the *Apodemus*/*Polyplax* studied here, which generally shows stronger degree of population structure in the parasite compared to the host both on the level of mtDNA (haplotype networks, AMOVA) and nuclear DNA (PCoA analyses).

#### Decoupled process of postglacial recolonization in host and parasite populations

Due to their intimate relationship, lice and their hosts share identical patterns of geographic expansion, unless the association is disrupted by a host switch. In other words, geographic distribution of a louse species/population is believed to be entirely determined by the host(s) (Marshall 1981). It is therefore remarkable that in our system we detected a decoupled process of recolonization by *A. flavicollis* and “its” specific lineage of *P. serrata*. As shown in the Figure 6, the two distinct mtDNA lineages of *A. flavicollis*, expanding from different refugia, are now distributed sympatrically across the whole sampled area and can be found on identical localities. Multilocus analyses show that this secondary postglacial encounter has been followed by frequent gene flow, resulting in a single panmictic population. In striking contrast, the two mtDNA haplotype clusters (S-W and S-E) of the *P. serrata* S lineage stopped their expansion from the glacial refugia at the narrow contact zone in the Central Europe (Fig 4). The inability of the two louse populations to cross the contact zone indicates that factors other than host-mediated distribution, or a mere within-refugia speciation, have played a role during the recolonization process. Based on the presented data, it is difficult to hypothesize on the probable cause of this discrepancy. However, an interesting possibility is presented by the symbiotic bacteria known to inhabit the lice ((Volf 1991; Hypša & Křížek 2007). The viability and/or reproduction of many blood feeding insects depend on various bacterial symbionts, and the intimacy of the host-symbiont association in such cases results in a metabolic cooperation between their genomes (Kirkness *et al.* 2010; Snyder & Rio 2013). The long-term isolation in refugium thus could lead to specific louse-genome vs. symbiont-genome adaptations that prevent an “incorrect” genome-genome combination.

In contrast to mtDNA, microsatellites did not show any apparent suture between the populations belonging to S-W and S-E subclades. While they clustered together geographically proximate populations, they did not provide information on higher hierarchical structure across Europe (Figs. 10 and 11). This picture is not surprising. Due to smaller effective population size and quicker coalescence compared to nuclear loci, mtDNA is considered to be a leading indicator of speciation processes (Zink & Barrowclough 2008). On the contrary, microsatellites should provide an appropriate tool for quantifying the volume of gene flow across the contact zone, after it is sampled more densely than in our current dataset.

#### Mitochondrial introgression from Polyplax lineage N to S

Despite the agreement between mtDNA and microsatellite in clustering the *Polyplax* samples into three major clades (N, S and Aa), a rare but striking discrepancy occurred in the CZLi population of *A. flavicollis*. Approximately half of the specimens sampled in 2005 (CZLi05N) clustered within the subclade A of the N lineage according to mtDNA, whereas microsatellites placed the sample within the S lineage (Fig. 8). The rest of the population sample (CZLi05S) was placed within the S lineage by both mtDNA and microsatellites Figs. 8 and 10). Such discrepancies are usually explained either by incomplete sorting of an ancestral polymorphism or by introgression after a secondary contact (Toews & Brelsford 2012; Hochkirch 2013). According to our view, recent mitochondrial introgression provides more probable explanation. We only found one instance across the whole Europe of shared haplotypes between distant louse lineages. The evolutionary age of the two mtDNA lineages is probably of pre-glacial origin (Hypša & Štefka 2008) making retention of ancestral polymorphism over such long period unlikely.

Recent population genomic studies revealed that species boundaries have not been as resistant to the gene flow of either mtDNA or nuclear DNA as previously thought (Harrison & Larson 2014). MtDNA introgression was detected especially in gender-biased gene flow (Zink & Barrowclough 2008). Coincidentally, parasites without free-living stages and intermediate hosts generally possess female-biased sex ratio (Criscione *et al.* 2005), which was confirmed also in *P. arvicanthis* lice from South African *Rhabdomys* (Mathee *et al*. 2007). Because the effective population size of mtDNA genes is four times lower than of autosomal genes, genetic drift influences mitochondrial haplotypes to a larger extent and can lead to faster fixation of unoriginal mitochondrial haplotypes (Funk & Omland 2003; Zink & Barrowclough 2008). Although mitochondrial introgressions occurring together with very low or even zero introgression of nuclear genes are rare, recent studies show that they occasionally happen, for example in Galapagos mocking birds (Nietlisbach *et al.* 2013) and North American chipmunks (Good *et al.* 2015).

Majority of studies on genetic introgression in animals analyse events of historical (pre- or postglacial) origin. In contrast, here we found that the introgression in *Polyplax* was probably very recent and short-lived, because repeated sampling at the locality in 2008 and 2014 did not reveal any introgressed haplotypes. Such dynamic development, where genetic information is quickly lost (or fixed), is in agreement with the biology of louse populations. Small, fragmented populations of lice are prone to rapid changes in their size and genetic composition.

#### Host specificity governs parasite dispersal and population size

The dispersal of parasites is to a great extent influenced by host sociality and vagility (Criscione *et al.* 2005; van Schaik *et al.* 2014; Mazé-Guilmo *et al.* 2016). Since parasitic lice inhabit a single host during the entire life cycle, their opportunities to spread are limited to direct host contact or to shared host shelters (Marhall 1981). Correspondingly, ectoparasite populations were recently shown to be more genetically fragmented than their hosts (Koop *et al.* 2014; Harper *et al.* 2015). A contradicting example was reported for *Polyplax arvicanthis* which maintains gene flow among populations of its four main *Rhabdomys* host species, perhaps due to the host’s interconnected distribution and relative sociality (du Toit *et al.* 2013). In our study system, genetic mixing of lice between the S and N lineages, was not observed even though both lineages share *A. flavicollis* as a common host species, possibly due to their older evolutionary origin and longer glacial separation. However, an interesting situation arose within the nonspecific N lineage of *Polyplax serrata*, where most subclades were not restricted to a single host species and microsatelite data did not show host-related genetic differentiation. Thus, occasional host switching between *A. flavicollis* and *A. sylvaticus* must occur. Since the two hosts generally avoid direct contact (Wasimuddin *et al.* 2016) and lice possess extremely short survival outside their host, lasting no longer than days (Marshall 1981), the switch is probably facilitated via shared burrows or scavenging.

When comparing dispersal activities of sucking lice and their hosts, one should expect higher level of historical gene flow in mice and lower level for *Polyplax* lice because of life history traits of the parasites (Kim 2006; Harper *et al.* 2015). Autocorrelation coefficient revealed markedly higher values for both *Polyplax* lineages compared with *Apodemus* hosts, especially over shorter distances, related to lower level of gene flow. Furtermore, high rate of *He* deficiency in louse populations (Table S2, Supporting information) indicates that gene flow is limited even within host populations, between lice from host individuals, which is in agreement with earlier results of Koop et al (2014) and Harper et al (2015). These findings support our expectations that host dispersal is the driving and limiting factor for parasite’s gene flow. Higher ability of dispersion in the more generalist lineage was also visible from IBD analyses of individuals, where Euclidian distances were significant for both S and N lineages, but with greater correlation coefficients in the S lineage (Fig. S13, Supporting information). These results agreed with Nadler‘s hypothesis predicting that specialists show greater IBD due to much fewer opportunities for finding suitable hosts in comparison with more generalist parasites like *Polyplax* from lineage N.

Another piece of evidence corroborating Nadler’s hypothesis was provided by direct comparison of genetic diversities between sympatric populations of the S and N lineages. In a global statistics of the whole lineages Fst index was significantly lower in the N lineage compared to S ((Table S3, Supporting information). These results indicated that lice from the S lineage (specialists) have smaller effective population sizes and more fragmented populations, which is associated with decreased frequency of heterozygotes. More importantly comparison of gene diversities between 7 sympatric pairs of N and S populations reached the same conclusions as the indexes calculated for the whole lineages (Fig. 14). Hence our unique system comprising multiple populations of two sister parasitic lineages provided clear evidence that even moderate shifts in host-specificities translate into significant differences in the genetic character of parasite populations.

In conclusion, considering the specific questions addressed by this study, we showed that: 1) the individual lineages of the parasite are well defined genetically (mtDNA and microsatellite differentiation) and by their affinity to different host species or different level of host specificity. However, contrary to general expectations, 2) genealogies of parasite lineages contained very little co-phylogeographical signal with their hosts. 3) We further revealed deeper population structure in parasites compared to their hosts, indicating limited dispersal in these parasites with single host life cycles. Finally and most importantly, 4) the patterns of diversity in two parasite lineages with differentdegree of host-specificity proved to follow the Nadler’s rule, i) stronger isolation by distance (resulting in deeper population structure) was seen in the single-host specific lineage and ii) substantial differences in genetic diversity were found between sympatric populations of the two lineages. In addition, we detected an unexpected geographical divison between two mitochondrial subclades of the parasite with lack of any discernible structure in the host populations. Our results indicate that the process of host-parasite co-evolution can adhere to predictable ecological phenomena (such as the link between host specificity and genetic diversity), but it also generates intriguing local patterns.

## Acknowledgements

We thank numerous colleagues from CZ and abroad, and students of the University of South Bohemia, CZ for their help with obtaining study material. This work was supported by the Czech Science Foundation (projects P505/12/1620, 14-07004S). Part of the work was carried out using resources of the National Grid Infrastructure MetaCentrum (project CESNET LM2015042).

## Data Accessibility

DNA sequences obtained in the frame of the study will be submitted to GenBank upon acceptance of the MS.

DNA alignments and microsatellite datasets were submitted to Dryad database (doi:10.5061/dryad.5jh39)

## Author Contributions

This study forms part of the PhD research of J.M., who performed laboratory and data analyses under the supervision of J.Š. V.H. and J.Š. conceived the study of *Apodemus*/*Polyplax* co-evolution. All three authors contributed toward the design of the study, and drafted the manuscript.

**Table S1:** Polymorphism in 16 loci amplified for *Polyplax serrata.*

| Locus name | Number of alleles in populations | Number of populations with polymorphic locus |
| --- | --- | --- |
| L1 | 1-6 | 25/32 |
| L2 | 1-10 | 32/32 |
| L3 | 1-5 | 22/32 |
| L4 | 2-11 | 32/32 |
| L5 | 1-7 | 28/32 |
| L6 | 1-5 | 23/32 |
| L7 | 1-8 | 32/32 |
| L8 | 1-10 | 28/32 |
| L9 | 1-7 | 31/32 |
| L10 | 1-3 | 16/32 |
| L11 | 1-9 | 31/32 |
| L12 | 1-6 | 28/32 |
| L13 | 1-8 | 32/32 |
| L14 | 1-11 | 29/32 |
| L15 | 1-11 | 28/32 |
| L16 | 1-11 | 31/32 |

**Table S2\***: Observed heterozygosity and HW deviation per each *Polyplax serrata* population and locus. Pop S – populations belonging to the S lineage; Pop N – populations belonging to the N lineage; levels of significance: ns – nonsignificant, * - P (0.05-0.01), ** - P (0.01-0.001), ** - P(<0.001). *\*provided as a separate file*.

**Table S3\***: Pairwise F_ST_ values between populations of *Polyplax serrata* S lineage, N lineage, *Apodemus flavicollis* and *A. sylvativus*. *\*provided as a separate file*.

**Table S4:**
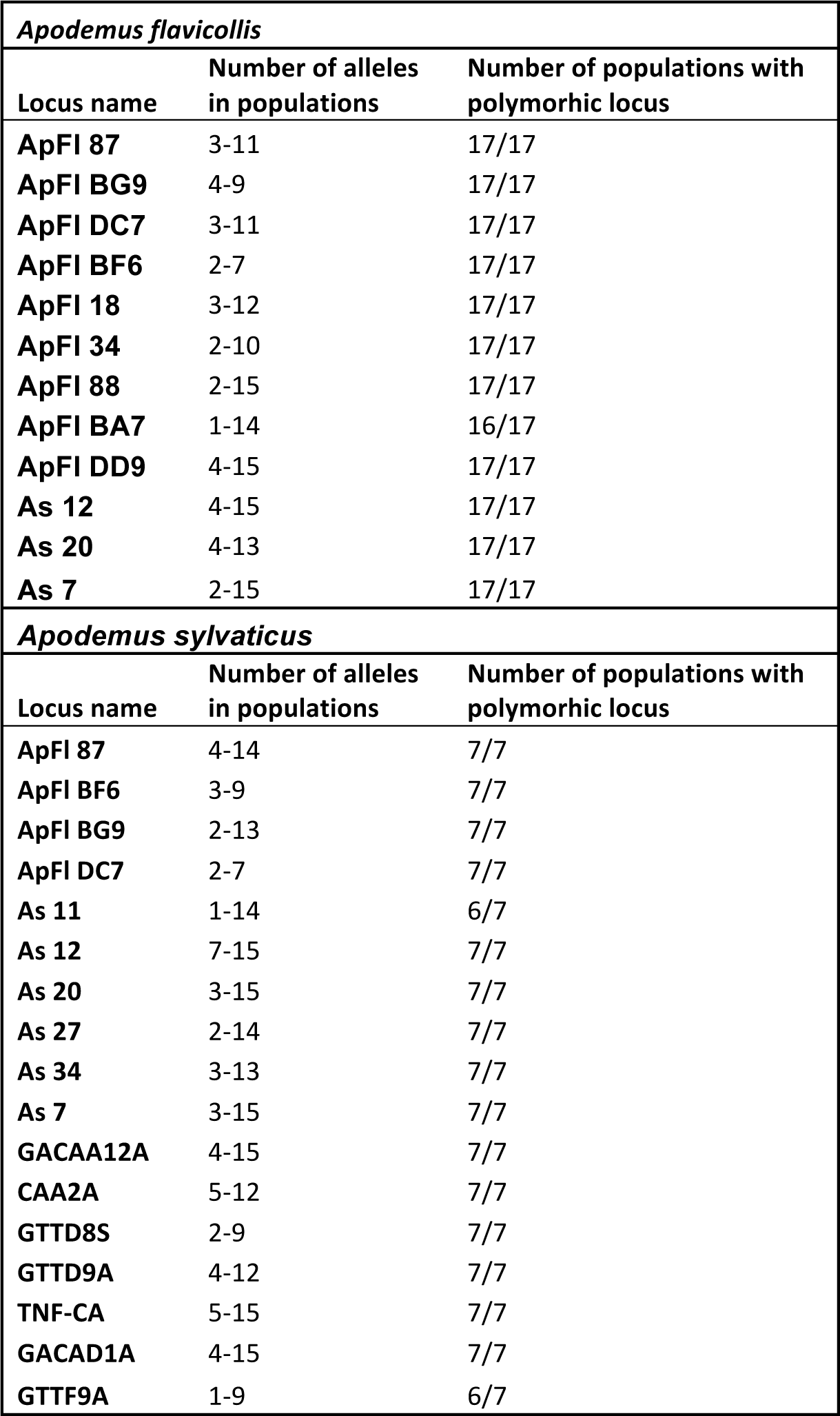
Polymorphism in microsatellite loci amplified for *Apodemus flavicollis* and *sylvaticus*.

**Table S5\***: HW deviation per each *Apodemus flavicollis* population and locus. Populations with less than 5 individuals were not analysed. Levels of significance: ns – nonsignificant, * - P (0.05-0.01), ** - P (0.01-0.001), ** - P(<0.001). *\*provided as a separate file*

**Table S6\***: HW deviation per each *Apodemus sylvaticus* population and locus. Populations with less than 5 individuals were not analysed. Levels of significance: ns – nonsignificant, * - P (0.05-0.01), ** - P (0.01-0.001), ** - P(<0.001). *\*provided as a separate file*

**Fig. S1:**
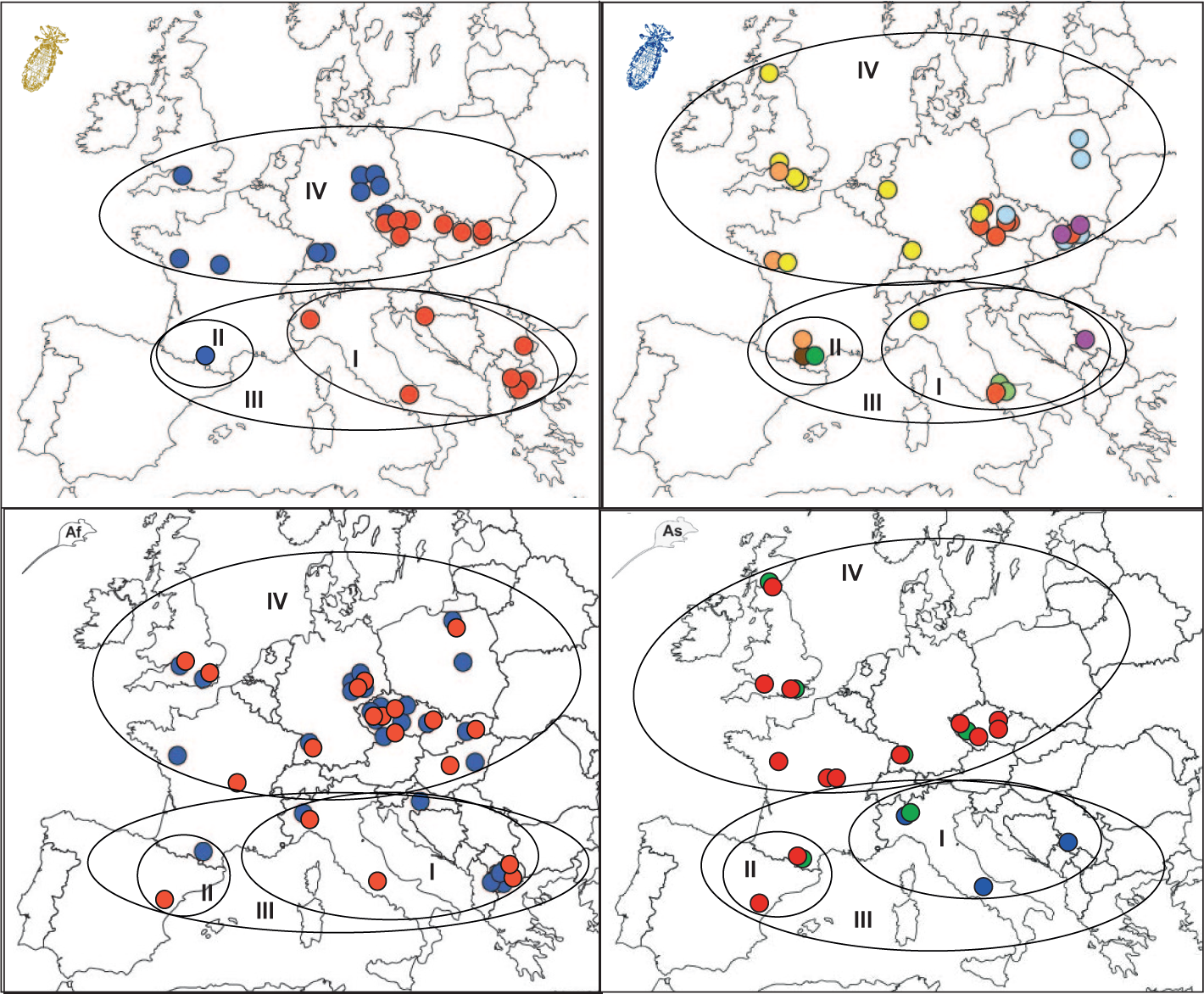
Hierarchical grouping of *Polyplax* populations (nonspecific and specific lineage separately), and *Apodemus flavicollis* (Af) and *A. sylvaticus* (As) into refugia and recolonized areas for DNASP and AMOVA analyses. Colour codes: as in Figs. 2, 6 and 7.

**Fig. S2:**
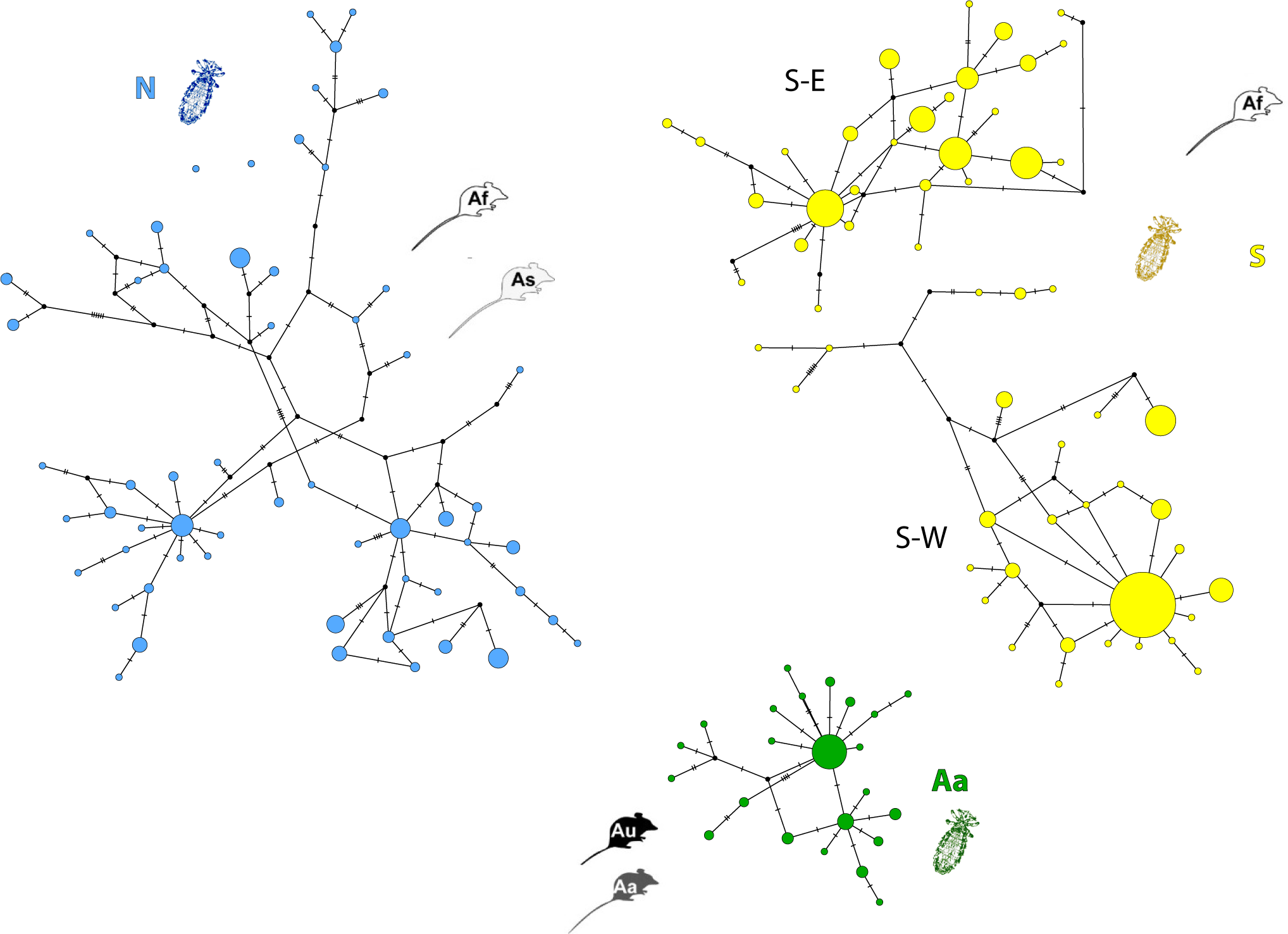
Haplotype networks for European lineages of Polyplax serrata. Networks were obtained in TCS program implemented in PopArt software using 381bp fragments of the COI gene. N – nonspecific lineage; S – specific lineage; Aa – lineage from *A. agrarius* and *uralensis*; host species abbreviations: Af – *Apodemus flavicollis*, As – *A. sylvaticus*, Aa – *A. agrarius*, Au – *Apodemus uralensis*; S-E and S-W – eastern and western clades of the S lineage, respectively.

**Fig. S3:**
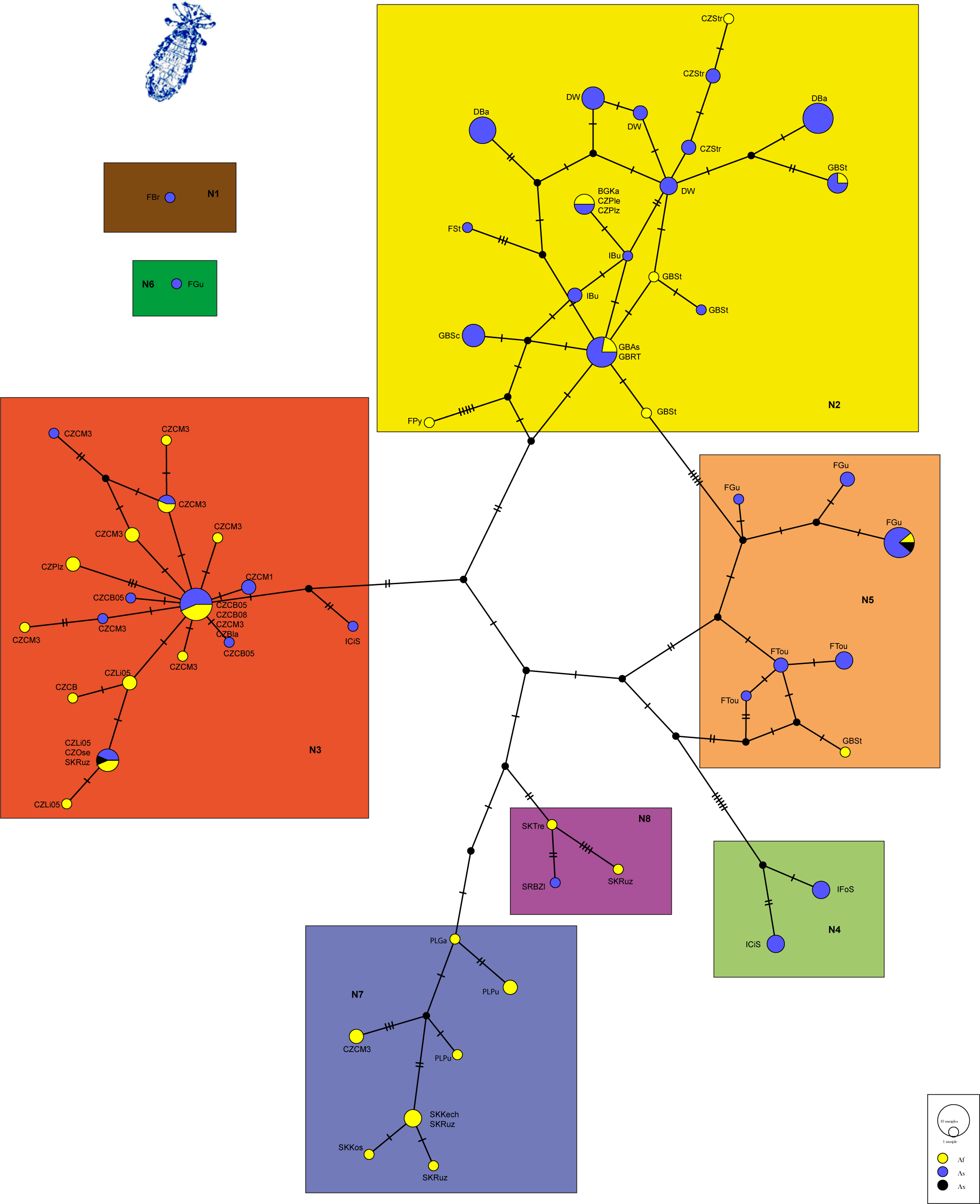
Haplotype network of the *Polyplax* N lineage with mitochondrial subclades and host species mapped. Network was obtained in TCS program implemented in PopArt software using 381bp fragments of the COI gene. Host species abbreviations as in Fig S1, Ax – host species not identified (either *A. flavicollis* or *A. sylvaticus*).

**Fig. S4:**
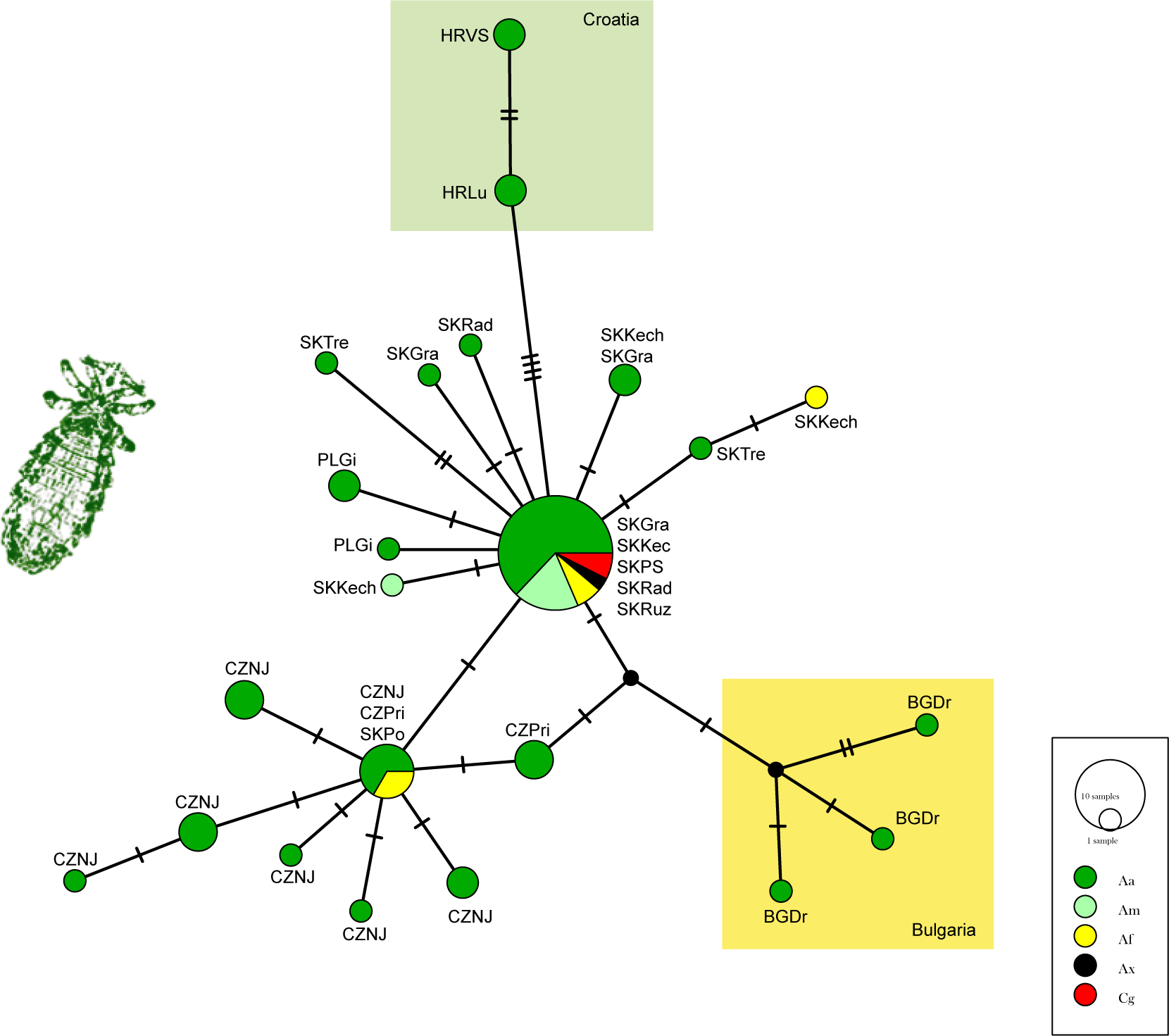
Haplotype network of the *Polyplax* Aa lineage with host species mapped. Distribution outside the Czech Republic and Slovakia highlighted in colour. Network was obtained in TCS program implemented in PopArt software using 381bp fragments of the COI gene. Host species abbreviations as in Figs. S1 and S2, Ax – host species not identified (either *A. flavicollis* or *A. sylvaticus*), Cg – *Clethrionomys glareolus*.

**Fig. S5:**
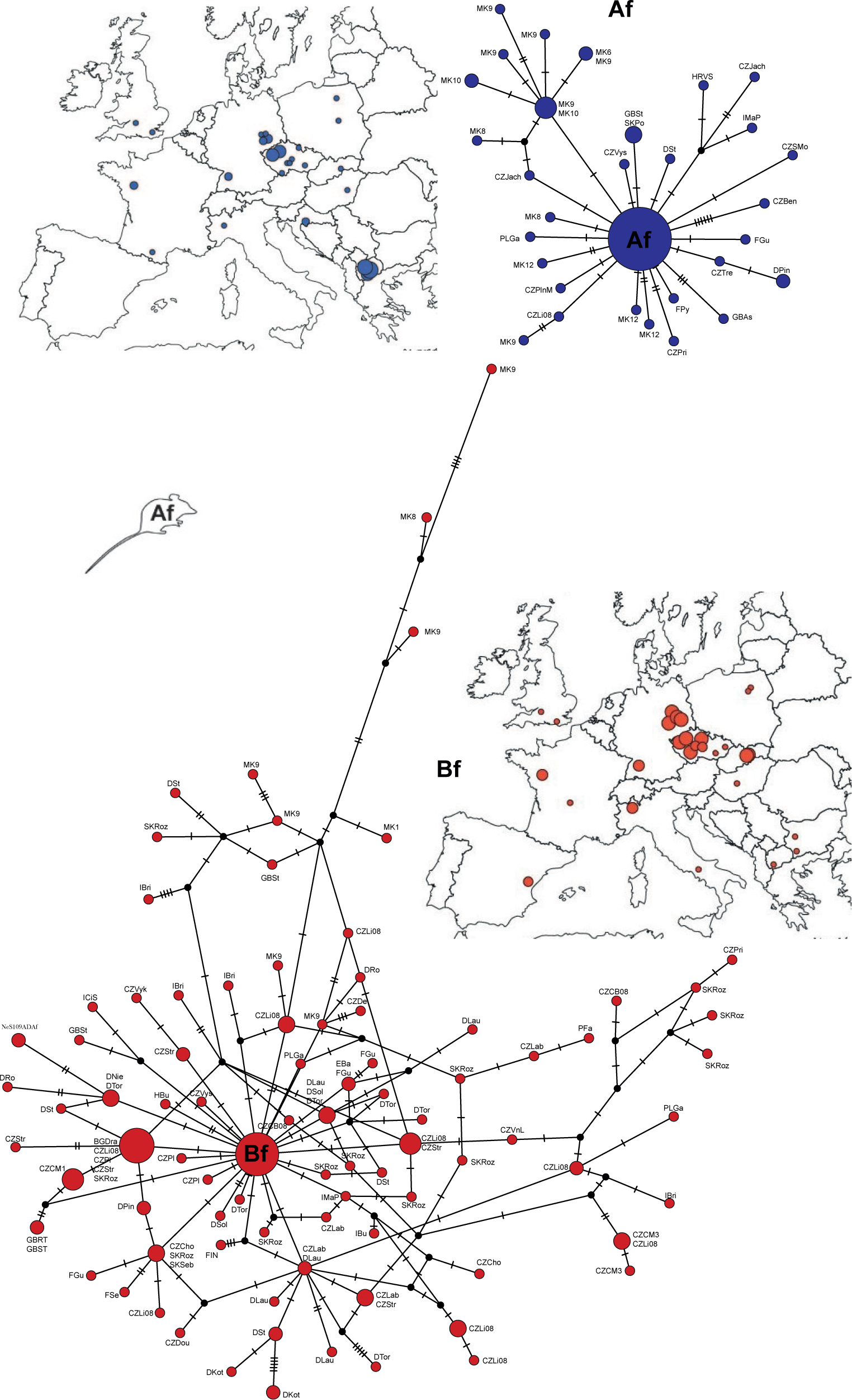
Haplotype networks and geographic distribution of the *Apodemus flavicollis* subclades Af and Bf. Networks were obtained in TCS program implemented in PopArt software using 1002 bp fragments of the mitochondrial D-loop.

**Fig. S6:**
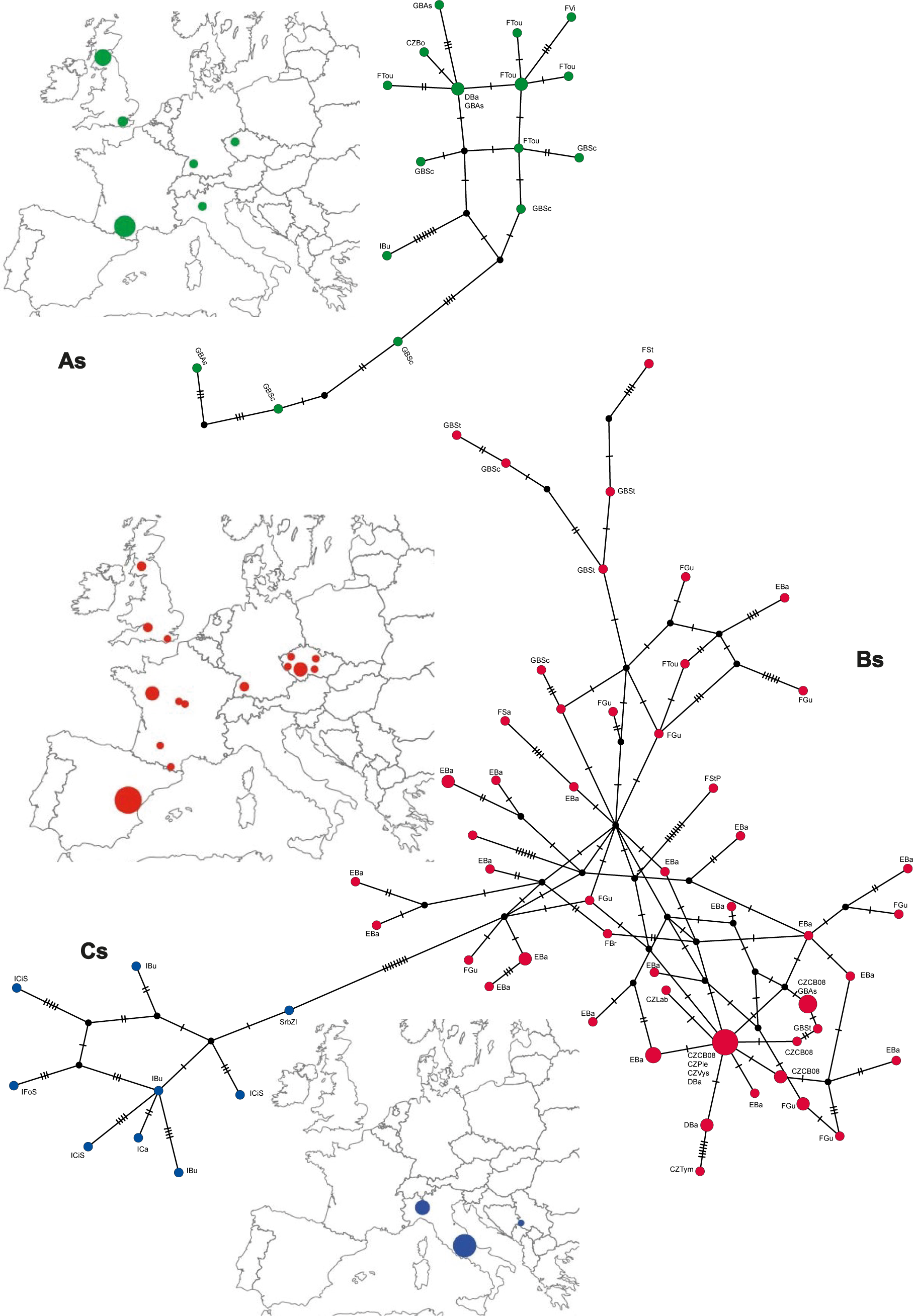
Haplotype networks and geographic distribution of the *Apodemus sylvaticus* subclades As, Bs and Cs. Networks were obtained in TCS program implemented in PopArt software using 1002 bp fragments of the mitochondrial D-loop.

**Fig. S7:**
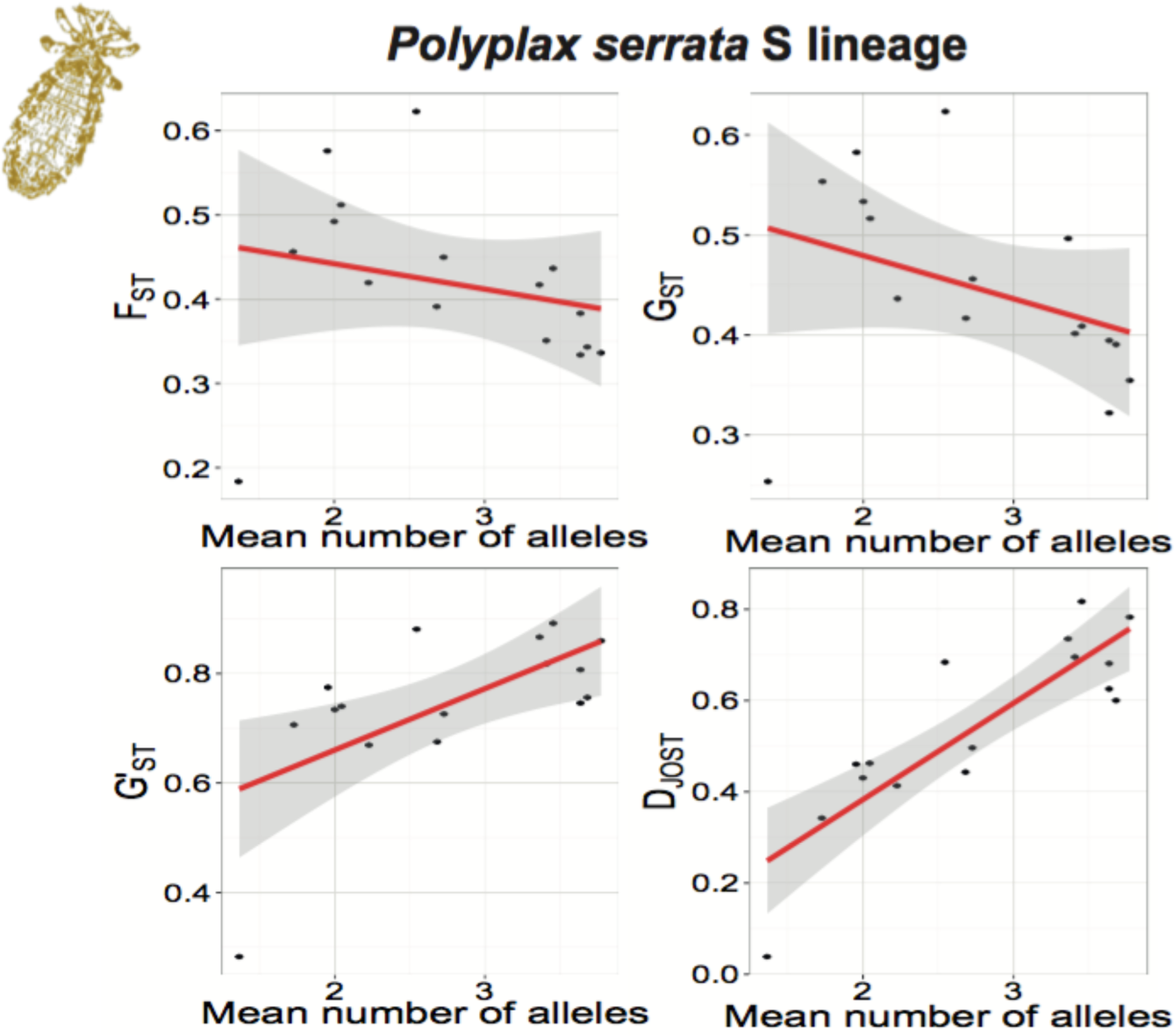
Interaction between FST statistics (and its derivates GST, G’ST and DJOST) and the mean number of shared alleles between *Polyplax serrata* populations of the S lineage.

**Fig. S8:**
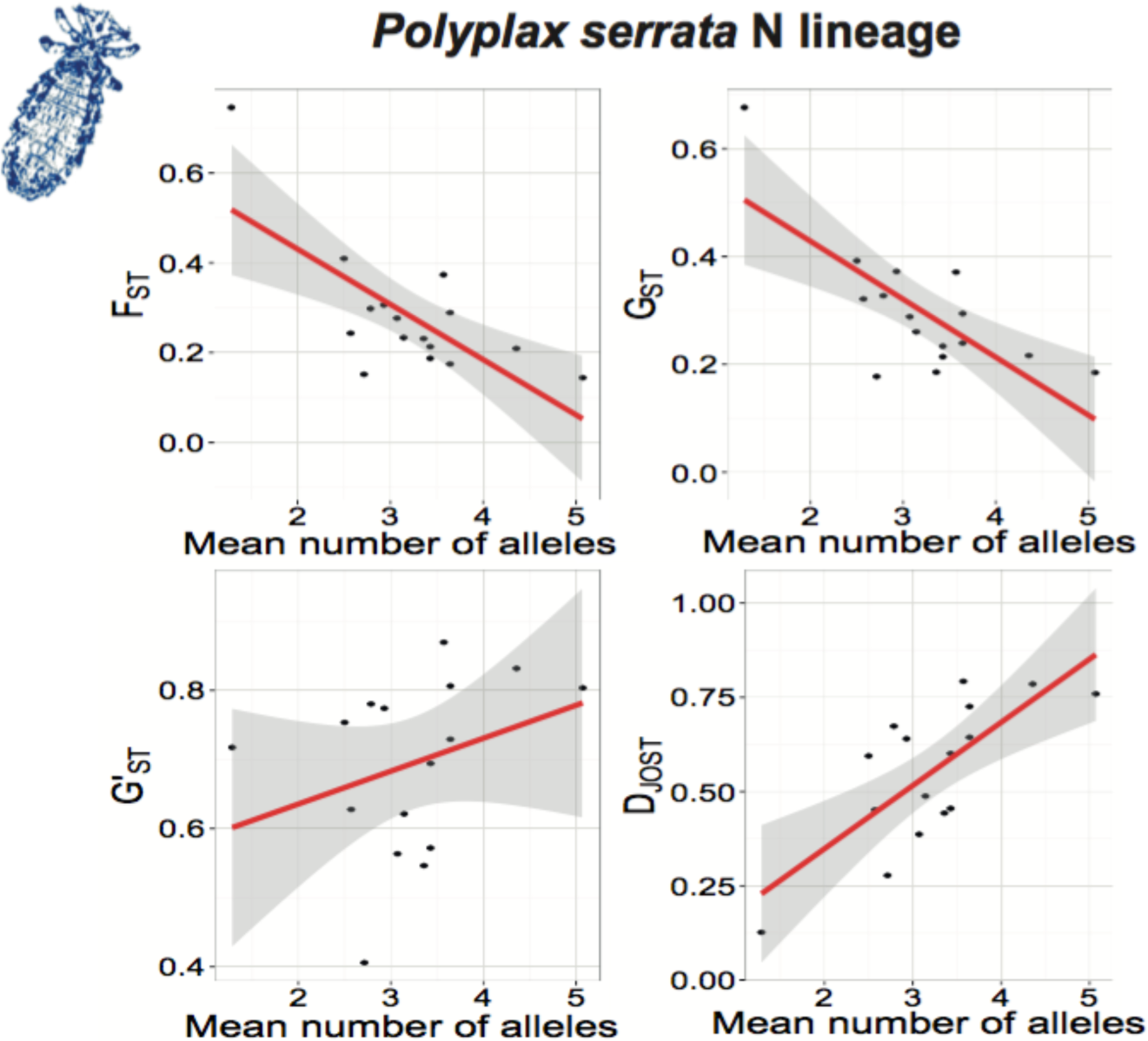
Interaction between FST statistics (and its derivates GST, G’ST and DJOST) and the mean number of shared alleles between *Polyplax serrata* populations of the N lineage.

**Fig. S9:**
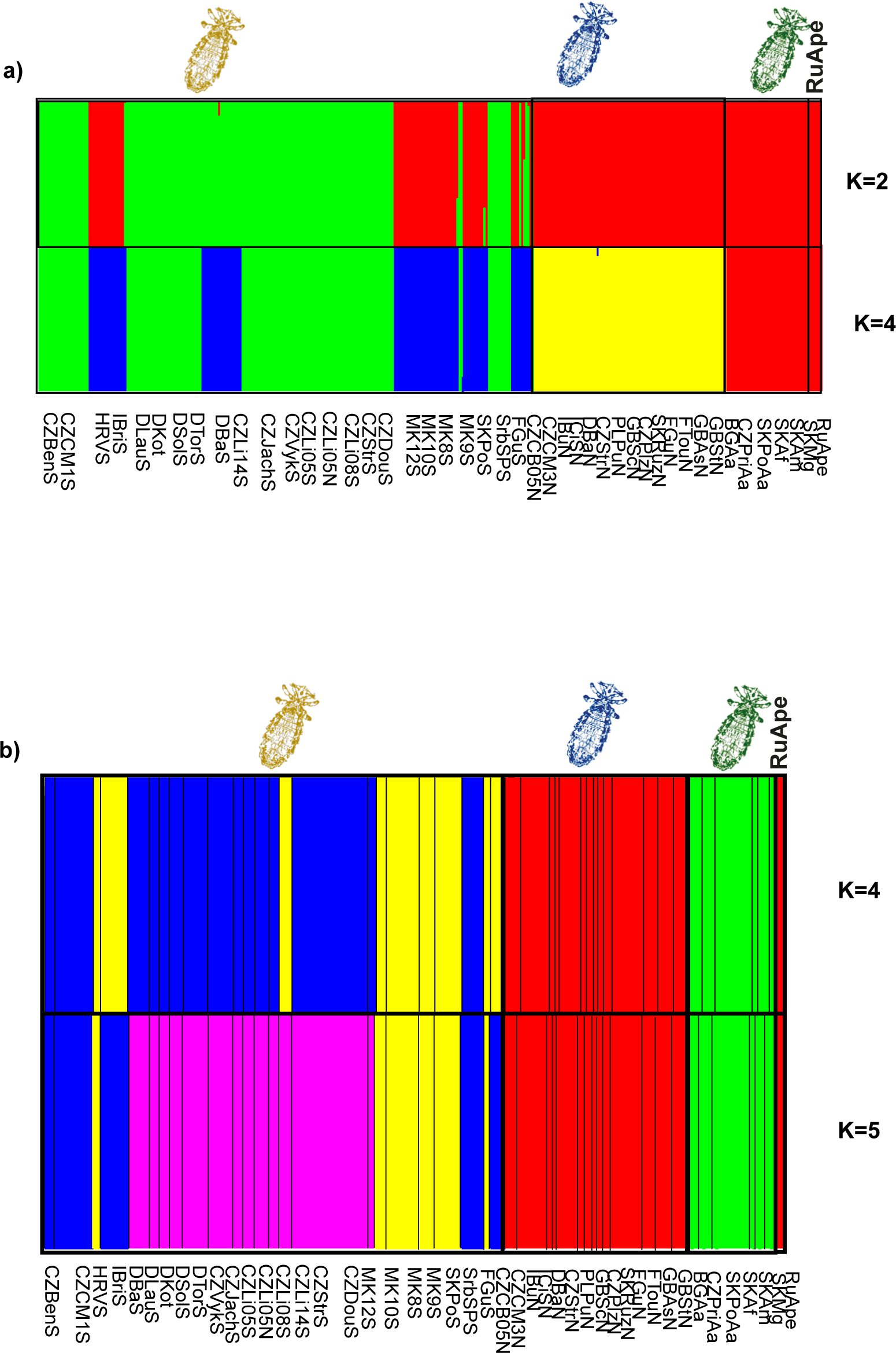
Bayesian clustering of *Polyplax serrata* individuals. a) Structure plots for K 2 and 4, b) BAPS plots for K4 and 5. Yellow louse image –mitochondrial S lineage, blue – N lineage, green – Aa lineage, Ape – ineage from Baikal Lake (*Apodemus peninsulae* host).

**Fig S10:**
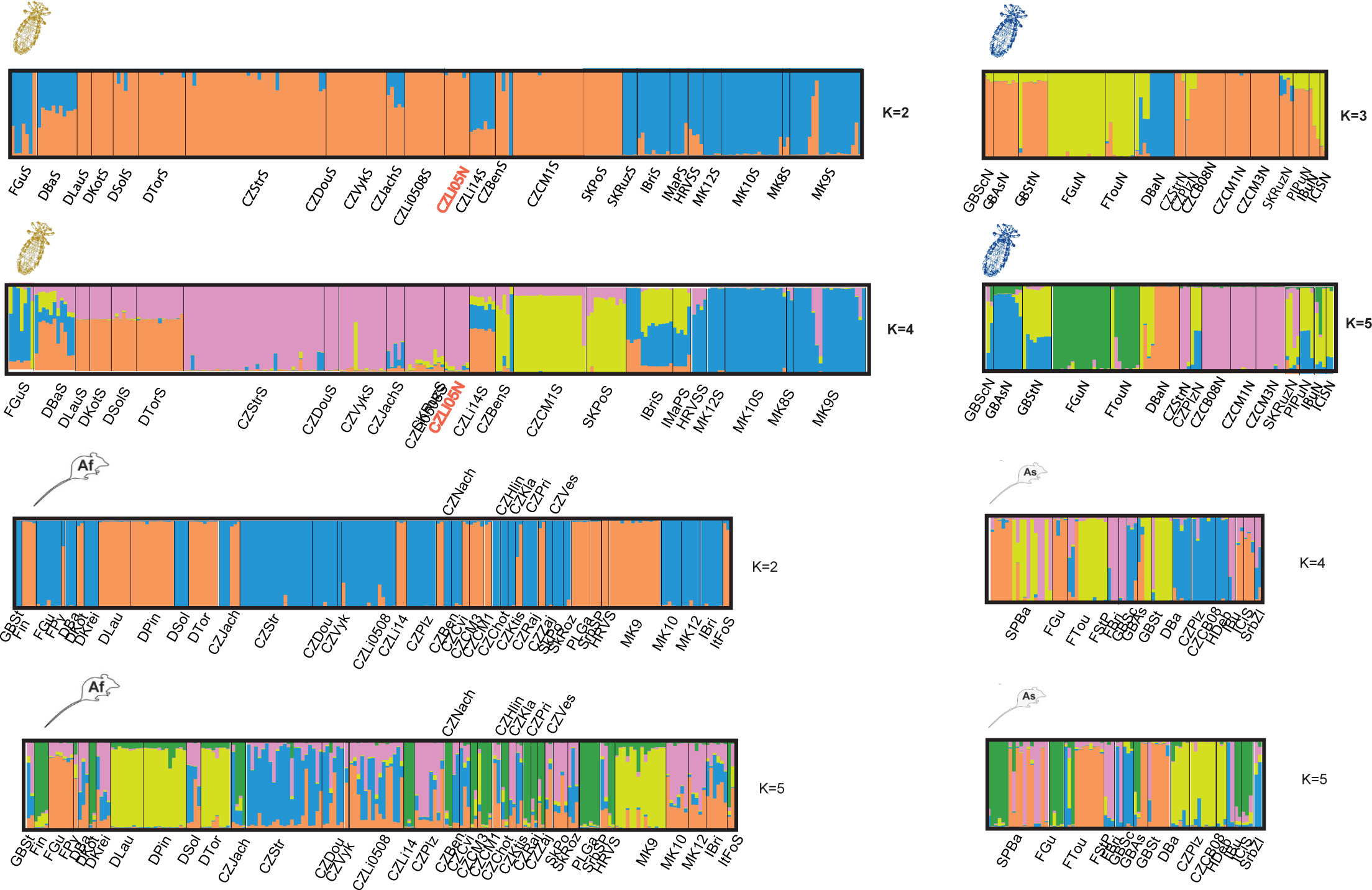
Structure plots for *Polyplax serrata* S lineage (yellow louse image, K2 and 4), *P. serrata* N lineage (K3 and 5), for *Apodemus flavicollis* (Af, K2 and 5) and *A. sylvaticus* (As, K4 and 5). Plots for K values represented by the two highest scores of ‘H are provided.

**Fig. S11:**
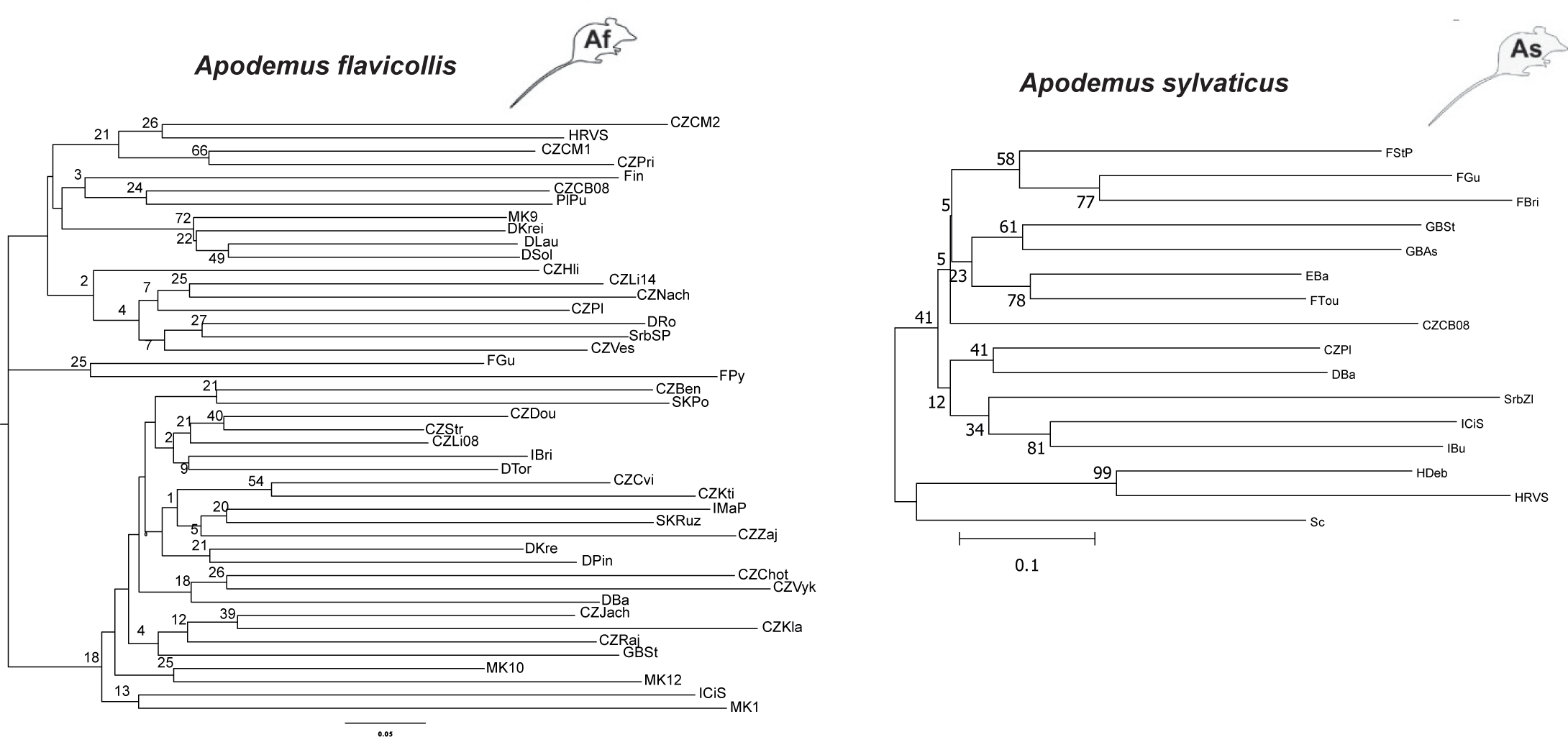
Neighbor-joining trees for *Apodemus flavicollis* and *sylvaticus* populations obtained in POPTREEW using pairwise DA values (Nei’s genetic distances) calculated from microsatellite data.

**Fig. S12:**
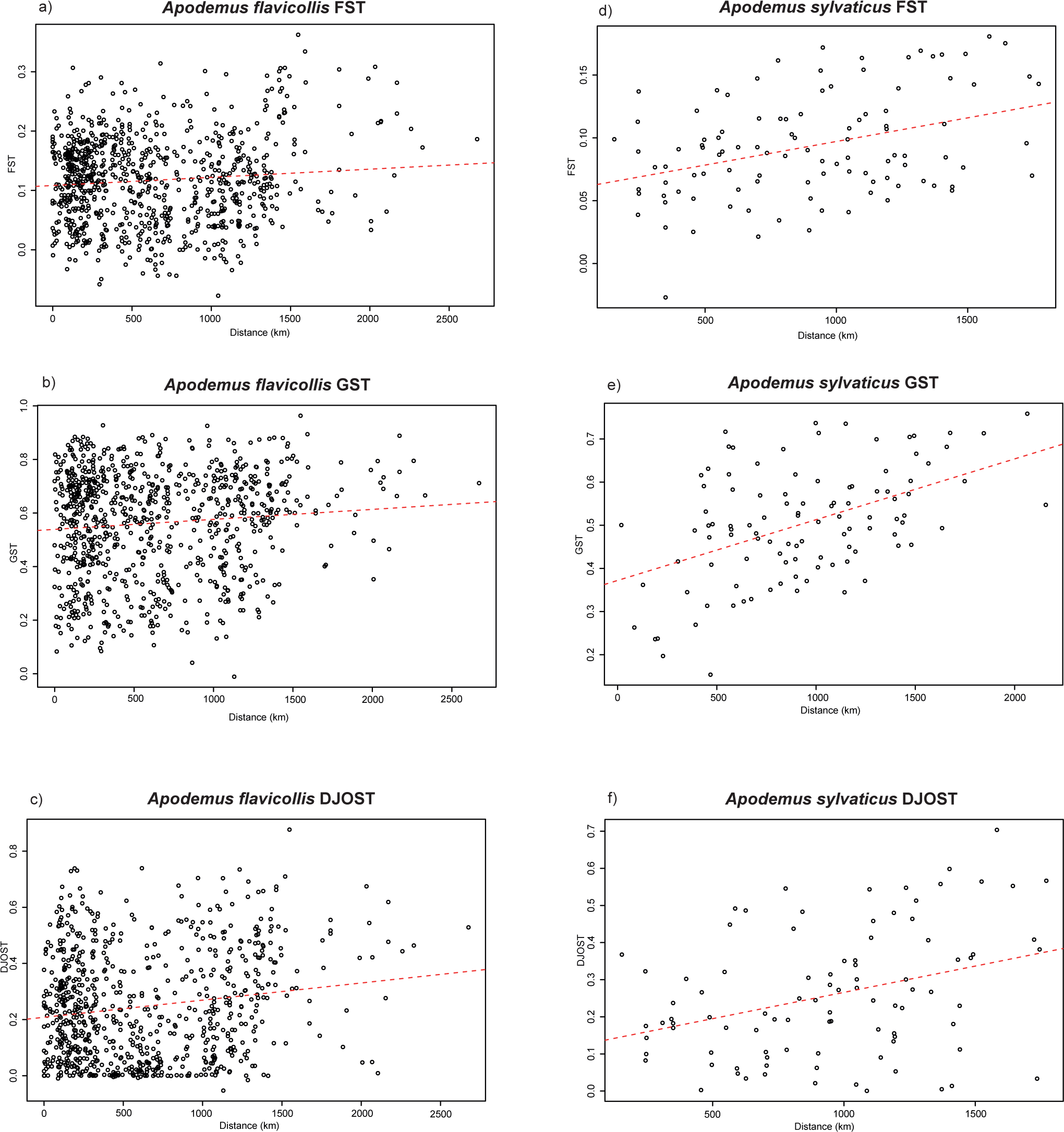
Mantel tests for correlation between genetic diversity indices (*F*_*ST*_, *G*_*ST*_ and *D*_*JOST*_) and geographic distance for the populations of *Apodemus flavicollis* (a, b and c) and *sylvaticus* (d, e and f).

**Fig. S13:**
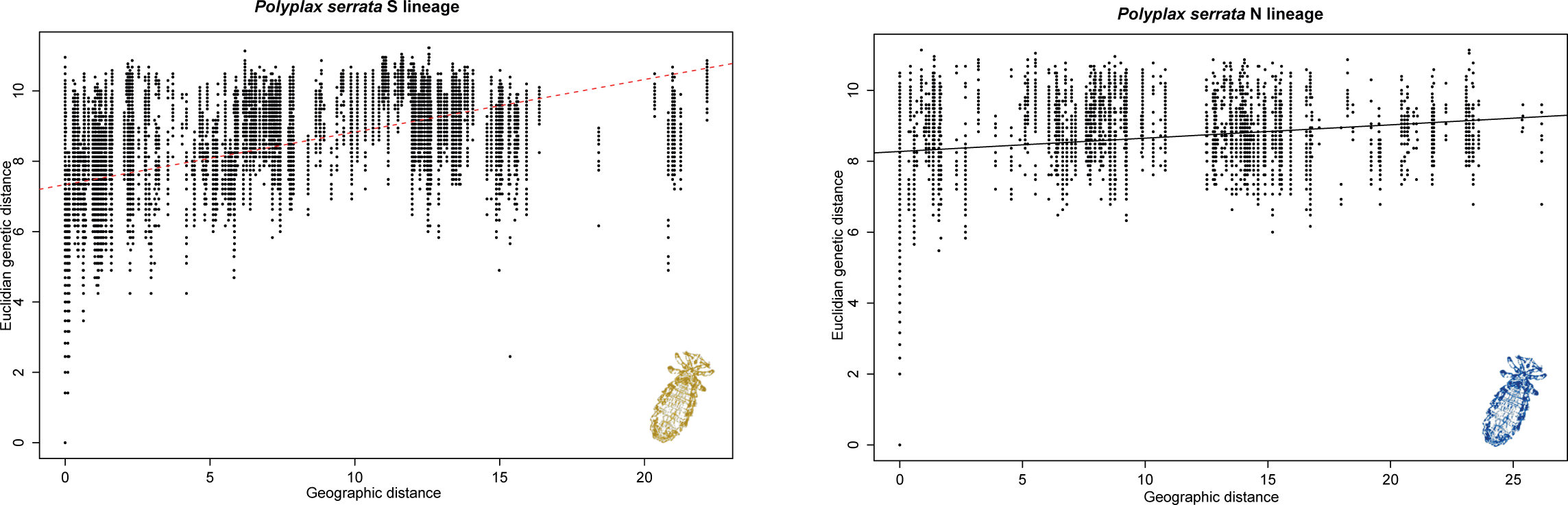
Correlation between Euclidean genetic distances and geographic distances for pairs of *Polyplax serrata* individuals. Plots were generated separately for S and N lineages in ADEGENET. Correlation was significant for the S lineage and non-significant for the N lineage (10 000 permutations).

**Document S1**: Supporting text providing information about PCR amplification, substitution models used in phylogenetic inference, and settings used in Bayesian clustering analysis of microsatellite data performed in STRUCTURE and BAPS programs.

## PCR amplification of mitochondrial genes

PCR reactions consisted of 1µL DNA, 1 µL of each primer (5µM), 10X High Fidelity PCR buffer, 2 µL MgCl2 (25mM), 2 µL dNTP Mix (2mM each), 0,2 µL High Fidelity PCR Enzyme Mix (5u/µL) (Thermo Fisher Scientific) and H2O added to a final volume of 20 µL. Thermal cycling conditions were held according to recommendation of the polymerase Enzyme Mix producer (Thermo Fisher Scientific) with initial denaturation for 3 min and 30 cycles of annealing temperature 50°C for primers L6625/H7005, 45°C for LCO1490/H7005 and temperatures recommended by Sweet et al. (2014) for nuclear genes. PCR products were purified with 0,2 µL of Exonuclease I (20u/µL) and SapI (10u/µL) enzymes each (New England Biolabs) and sent to Macrogen Europe (Macrogen Inc.) for sequencing.

PCR reactions had the same composition as for lice, Taq DNA polymerase (Top-Bio, Czech Republic) was used instead of the High Fidelity PCR Enzyme Mix. Thermal cycles started with an initial denaturation at 94°C 2 min, followed by 30 cycles of denaturation at 94°C for 15s, annealing at 49.5°C for primers 1/2bis, and 55°C for primers 3 and 4, elongation at 72°C for 1min, with a final extension at 72°C for 5min. Products were purified enzymatically as described above and sent to Macrogen Europe (Macrogen Inc.) for sequencing.

## Substitution models used in phylogenetic analyses

For Polyplax serrata COI 381bp dataset GTR+G model was used, for P. serrata concatenated alignment (COI and three nuclear genes) TRN+I+G was selected, for A. flavicollis and A. sylvaticus GTR+I+G and TN93+I, respectively, had the highest score.

## Structure and BAPS analyses

### Polyplax

In Structure, five independent analyses were run for K=1-6, separately under both admixture and no-admixture priors with 5 Million MCMC and a burnin of 500000. BAPS was run with the clustering of individuals method for K=2–6, re-analysed ten times for each number of the clusters followed by admixture estimation based on mixture clustering. The H‘ statistic from CLUMPP 1.1.2 (Jakobsson & Rosenberg 2007) that measures similarity of runs was used when determining the optimum K value.

#### Apodemus

Analyses were run as above with the number of clusters in Structure K=1-5, 1 Million MCMC, 100 000 burn-in and 10 independent runs.

